# AnnZarro: scalable, interactive exploration of cell-by-cell and gene-by-gene relationships in single-cell data

**DOI:** 10.64898/2026.10.10.758158

**Authors:** Dominik J. Otto, Siddharth Baasri, Manu Setty

**Author notes:** Corresponding authors: Dominik J. Otto, Manu Setty.

## Abstract

Cell-by-cell and gene-by-gene relationships, such as kernels and correlations, shape most single-cell analyses but are rarely inspected directly due to a lack of visualization tools that display them. AnnZarro is a point-and-click browser viewer that shows these matrices linked to an AnnData object’s cells × genes data. Using AnnZarro, experimental biologists can explore data without code and computational biologists can probe the relationships behind their analyses. One focused cell and one focused gene drive every panel. Clicking a cell paints its row of any cell-by-cell matrix over an embedding or spatial coordinates; clicking a gene paints its row of any gene-by-gene matrix over a gene plot such as a volcano. The focus also links genes to the cells where they change, and cells to the genes that distinguish them. AnnZarro reads only the matrix rows and columns each panel displays from zarr stores and hence users can explore matrices far larger than memory on a laptop or a shared server. Large datasets open on a reproducible subset, and users step through the remaining cells in parts. On a laptop, a 100,000-cell part of Tahoe-100M opens in 4 s and stays interactive. With the subset switched off, AnnZarro draws all 95.6 million cells in one plot in about 5 to 7 s. This tutorial prepares a dataset with Kompot differential abundance and expression, a diffusion kernel and gene correlations, then explores it in six worked examples. AnnZarro is available at https://annzarro.readthedocs.io.

## Introduction

Two of the most informative objects in a single-cell analysis are how similar each cell is to every other cell and how similar each gene is to every other gene. Both are computed routinely, as kNN graphs, diffusion kernels, distances and gene-gene correlations, and both drive downstream results such as trajectories, fate probabilities and gene programs. Explored directly, they answer questions that a summary cannot: which cells a given cell resembles in the space where its pseudotime and fate probabilities were computed, which genes change together with a gene of interest and how strongly, and how both answers shift from one cell state to the next. Static plots show one gene or one cell per figure, and existing interactive viewers rarely display these pairwise matrices. Of those we examined, only one colors an embedding by a cells × cells row, and only when its index is typed in. Users cannot view genes × genes matrices in any of them, or select a cell or gene in one plot to set the row shown in the others (**Fig. 1c**).

**Figure 1:**
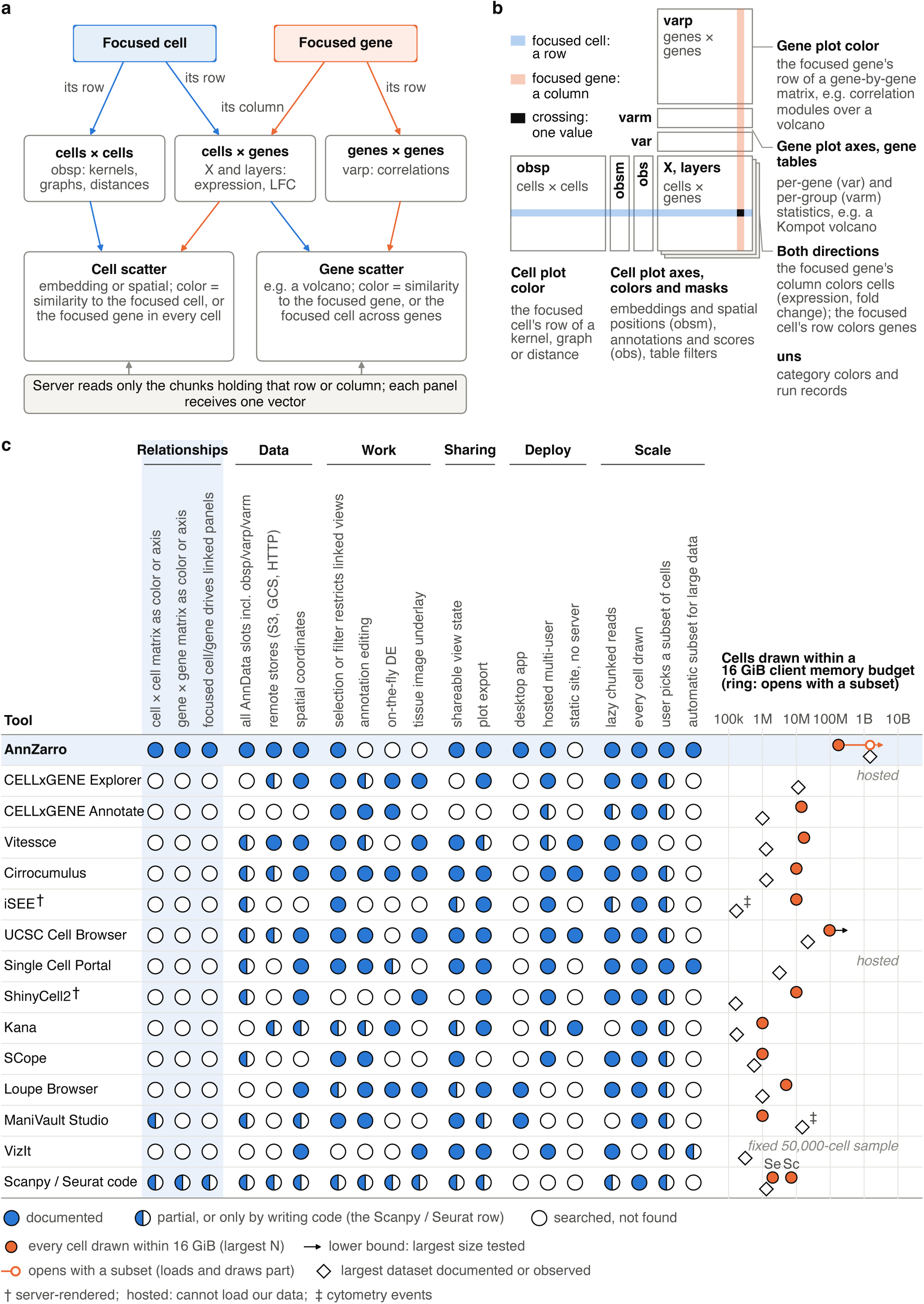
The focus model, the AnnData slots behind it, and AnnZarro compared with other viewers. **a**, The focused cell selects rows of the cells × cells and cells × genes matrices (blue); the focused gene selects a column of cells × genes and a row of genes × genes (orange). Each scatter is colored by the selected vector, and clicking a point moves the focus. LFC, log fold change. Interactive walkthrough: https://annzarro.readthedocs.io/en/latest/tutorials/tour.html. **b**, Which stored array drives which view, on an AnnData object laid out around its cells × genes matrix. A click on a cell selects one row (blue) of every cell-indexed array; a click on a gene selects one column (orange) of every gene-indexed array. AnnZarro fetches only these slices; their crossing in X or a layer is the focused cell’s value for the focused gene. Rows of obsp color cell plots, rows of varp color gene plots, and X and layers link cells and genes in both directions. **c**, Feature comparison. Rows are tools, columns capabilities grouped by job. CELLxGENE is shown as its two programs; Loupe Browser 9.1.0 was checked in the running app on demo data, with spatial and image underlays from its documentation and lazy reads measured in the app (**Supplementary Note 1**); Scanpy / Seurat is the baseline of what code can do. Filled circle, documented and working in the GUI; half-filled, partial or only by writing code; open circle, searched and not found. In the shaded Relationships columns, only AnnZarro reads genes × genes matrices and lets a focused cell or gene choose the row that every linked panel shows. AnnZarro lacks annotation editing, on-the-fly differential expression, tissue image underlays and a static-site mode. The Cells strip (log scale, 100,000 to 10 billion cells) shows, as a filled orange circle, the largest dataset built from Tahoe-100M of which each tool drew every cell without failing and within a 16 GiB client memory budget († server-rendered: its server is measured instead); what stopped each tool at the next size (client or server memory, a crash, an unreadable input) is given in **Supplementary Note 1**. Ring, the largest store drawn with a subset; open diamond, the largest dataset the project documents (for AnnZarro, the largest store tested here, with a subset); arrow, a lower bound at the largest size tested; ‡, cytometry events. Sc, Scanpy; Se, Seurat. Method, rules and sources are in **Supplementary Note 1**.

Similarity is only one kind of cells × cells matrix. In spatial transcriptomics, neighbor graphs and physical distances between cells are stored as such matrices ^1^, and so is cell-to-cell communication inferred under the constraint of how far a ligand can diffuse through the tissue ^2^. In the nervous system, a connectome is a matrix of synapses between individual neurons, which can now be read against the neurons’ transcriptomic types ^3,4^. Each of these matrices answers a question about one cell at a time: which cells it neighbors, signals to or is wired to. A static figure answers such a question for one cell. Asking it for many cells in turn, and for the genes that set them apart, requires a view that redraws with each click.

We built AnnZarro to make these matrices explorable. AnnZarro uses AnnData objects ^5^, the format that scanpy and the scverse tools write, where cells × cells matrices are stored in obsp and genes × genes matrices in varp, beside the cells × genes data in X and layers (**Fig. 1a,b**). Clicking a cell paints its row of a cell-by-cell matrix over any embedding or over the positions of cells in a tissue section. Results computed in the original space, such as diffusion-based pseudotime, can be checked against the matrix that produced them. Clicking a gene paints its row of a gene-by-gene matrix over any gene-level plot, such as a differential expression volcano. The same click links cells and genes in both directions: a focused gene colors cells by its expression or fold change, and a focused cell colors genes by its values. This link matters most for differential expression methods that estimate a fold change for every cell, such as Kompot ^6^. Both the matrices and the datasets are large: a cells × cells kernel over 100,000 cells holds 10^10^ values, and single-cell datasets now reach nearly 100 million cells ^7^. AnnZarro reads only the matrix rows and columns each panel displays, and on a laptop users can browse all 95.6 million cells of Tahoe-100M, through a reproducible subset or in one plot (**Fig. 1c**).

### Development of the protocol

AnnZarro was developed to inspect the outputs of Palantir ^8^, Mellon ^9^ and Kompot ^6^, which store their results across every AnnData slot, including dense cells × genes layers and cells × cells kernels that the existing viewers could not display. It builds on two ideas from other groups: panels linked so that a selection in one drives the others, as in iSEE ^10^, and views that are shared as configurations, as in Vitessce ^11^. To these it adds a focused cell and a focused gene that select rows of stored pairwise matrices, and lazy reads of chunked zarr stores so that these matrices need not fit in memory. Its design rests on five principles.

1. **A focused cell and a focused gene.** The application keeps one focused cell and one focused gene, with back and forward history. Clicking a point in any panel moves the focus. Every panel whose content depends on the focus fetches the new slice and redraws in place, keeping its zoom.
2. **Any data on any axis.** A cell scatter can take the x, y and optional z positions of its points, and their color, from any per-cell value. These include cell metadata in obs, a column of any obsm matrix (such as an embedding or spatial positions), a gene’s column of X or any layer (such as expression, imputed expression or fold changes), and the focused cell’s row of any cells × cells matrix in obsp. A gene scatter mirrors this with gene metadata in var, a column of any varm matrix, the focused cell’s row of a layer and the focused gene’s row of any genes × genes matrix in varp. Several cell and gene scatters can be open at once, and a click that changes the focused cell or gene redraws every panel that depends on it. Tissue positions are one more obsm matrix, so spatial data need no special mode. Coloring a tissue section by the focused cell’s row of an expression kernel shows where similar cells lie. Tables of cells or genes define masks with arbitrary AND/OR conditions, and plots follow them.
3. **Bounded memory on both sides.** AnnZarro is built for dense data: imputed and fold-change layers, dense kernels and correlation matrices, which are too large to load but are stored in chunks. The server opens zarr stores lazily and reads only the chunks a request touches, so one gene of a dense layer or one row of a dense kernel costs one chunk column or row. The browser receives only the vectors on screen and, by default, for datasets of more than 200,000 cells only their values for a reproducible subset, whose parts reach every cell. Neither side holds the full dataset, so a laptop can explore data that live on a cluster.
4. **Read-only hosting.** The server never writes to datasets and runs no user code. Datasets on institutional servers can be exposed, with login, without granting write or compute access.
5. **Views as shareable objects.** Panel sets are stored on the server, so a view defined by one user can be opened by every other user of the instance. Deep links encode layout and focus in the URL, and plots export as SVG, PNG, JPEG or WebP.

The same code runs as a multi-user server, as a local server, or as a self-contained desktop application for macOS, Windows and Linux. **Fig. 2** shows the interface on the data used in this protocol.

**Figure 2:**
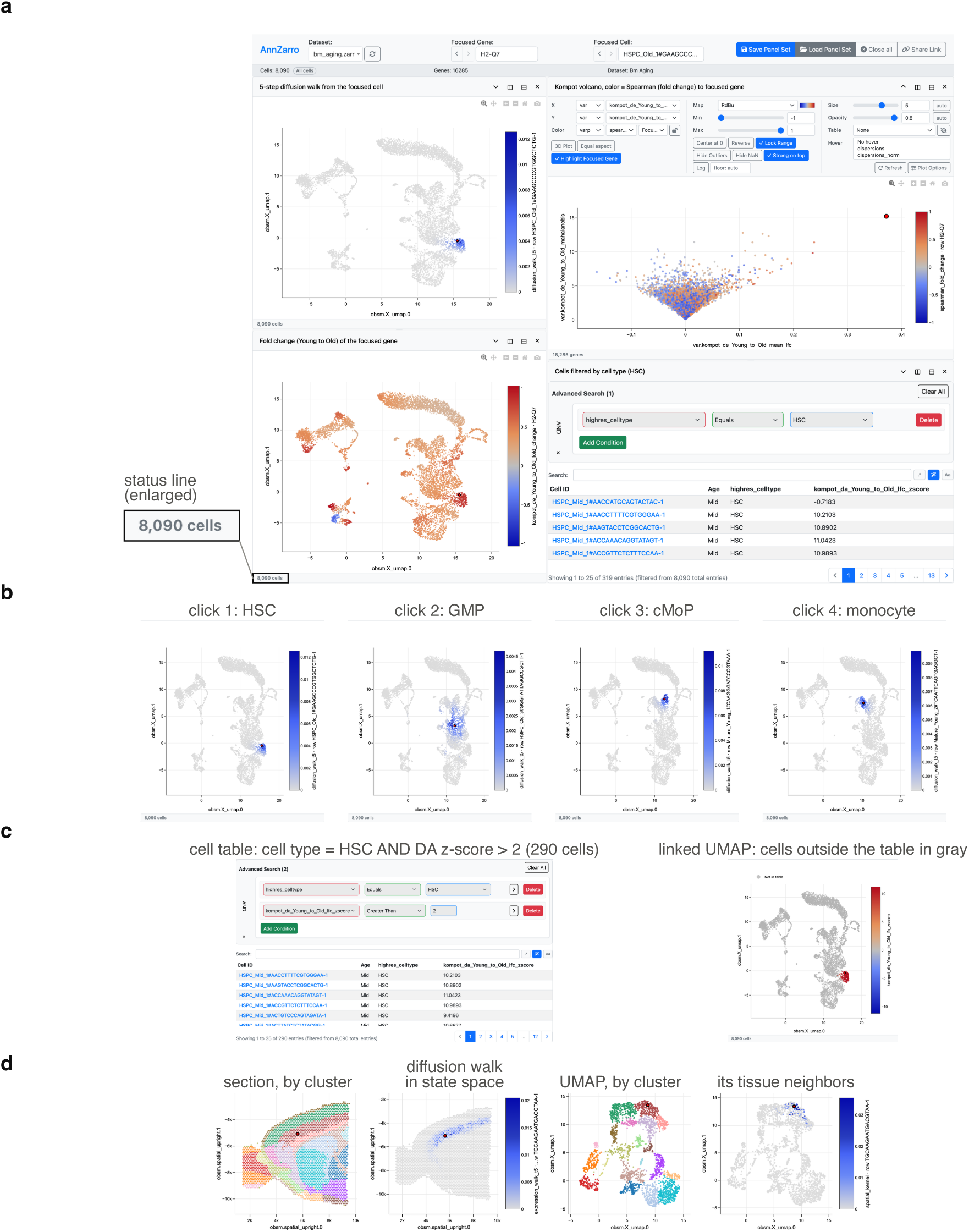
AnnZarro on the demonstration data. **a**, Four linked panels with the focused gene *H2-Q7* (histocompatibility 2, Q region locus 7) and a focused hematopoietic stem cell (header): the UMAP colored by the stem cell’s row of a five-step diffusion walk; the Kompot volcano colored by each gene’s fold-change correlation with *H2-Q7*; the UMAP colored by *H2-Q7*’s fold change; and a cell table filtered to stem cells. The volcano’s controls are open, with *Strong on top* pressed (the strongest values drawn last); the line under each plot is its status line (Step 34; inset, enlarged). **b**, The diffusion-walk panel after four successive clicks from a stem cell through a granulocyte-monocyte progenitor (GMP) and a common monocyte progenitor (cMoP) to a monocyte; each click fetched one row. **c**, A table filter combining two conditions (cell type is HSC and differential abundance z-score above 2; 290 of 8,090 cells) masks a linked UMAP: cells outside the table turn gray without any new data request. **d**, Mouse brain section profiled using Visium ^12^ after a click on a cortex spot: the section by cluster; the spot’s row of a diffusion walk in expression state space, which stays in its cortical layer without using tissue positions (88.7% of its mass falls in the spot’s Leiden cluster); the UMAP by cluster and by the spot’s row of a tissue-distance kernel.

### Overview of the procedure

The procedure takes users from a processed AnnData object to a set of linked, shareable views in five stages (**Fig. 3**).

1. **Installation (Steps 1–2).** Install the Python package or the desktop app.
2. **Data preparation (Steps 3–10).** Run Kompot differential abundance and expression on the Kompot tutorial dataset of aging murine bone marrow, add a diffusion-walk kernel to obsp and gene-gene Spearman correlations to varp, and write a chunked zarr store.
3. **Starting AnnZarro (Steps 11–13).** Start the server, open the store and add panels.
4. **Worked examples (Steps 14–36).** (1) Walk a diffusion neighborhood across the embedding (Steps 14–17); (2) browse co-regulated genes in a differential expression volcano (Steps 18–21); (3) go from a gene to the cells where it changes (Steps 22–24); (4) compare two cell states gene by gene (Steps 25–27); (5) mask plots with table filters (Steps 28–31); (6) browse a dataset of millions of cells, Tahoe-100M (Steps 32–36).
5. **Saving, sharing and hosting (Steps 37–44).** Save panel sets, share links, export figures and, optionally, host datasets for a lab.

**Figure 3:**
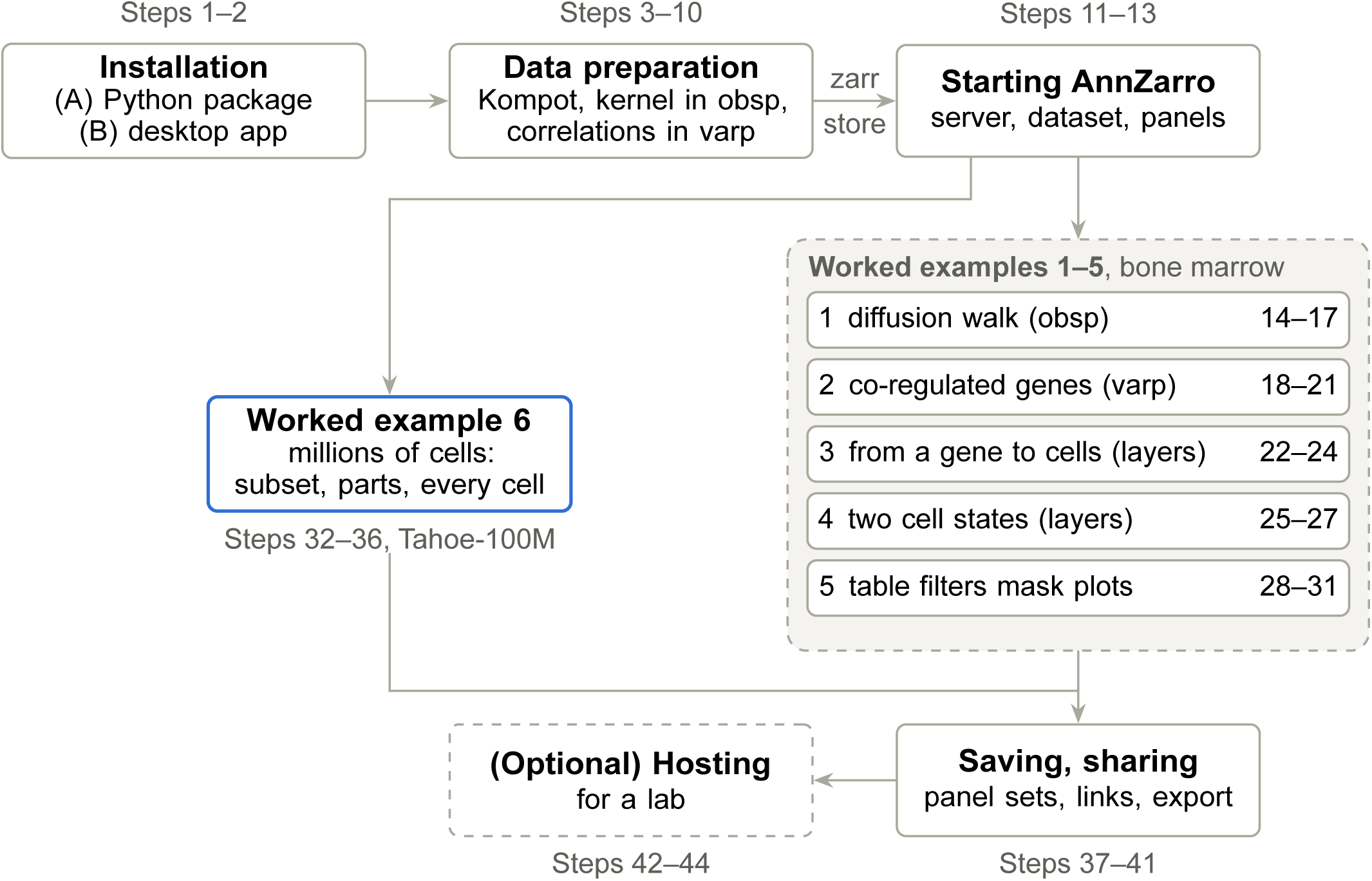
Overview of the procedure. Stages with their step numbers. Worked examples 1–5 use the bone marrow store prepared in Steps 3–10 and can be run in any order, except that Worked example 3 reuses the volcano of Worked example 2; Worked example 6 uses Tahoe-100M. Dashed box, optional stage.

Worked examples 1, 2, 4 and 5 can be run in any order after Step 13; Worked example 3 reuses the volcano of Worked example 2, and Worked example 6 uses a separate, large dataset.

### Applications

AnnZarro is built for results that live in matrices too large to plot at once: per-cell, per-gene and pairwise quantities that users want to browse one slice at a time. We have used it to explore the results of Kompot ^6^ as the method was being developed. Three classes of question motivate the protocol and each is demonstrated in it on the bone marrow data (**Figs. 4–6**), a connectome of *C. elegans* neuron classes shows the neuronal case (**Fig. 4d**) and a public Visium dataset of the mouse brain shows the spatial case, with the spots’ tissue positions as the axes and a spatial kernel in obsp (**Fig. 2d**).

**Figure 4:**
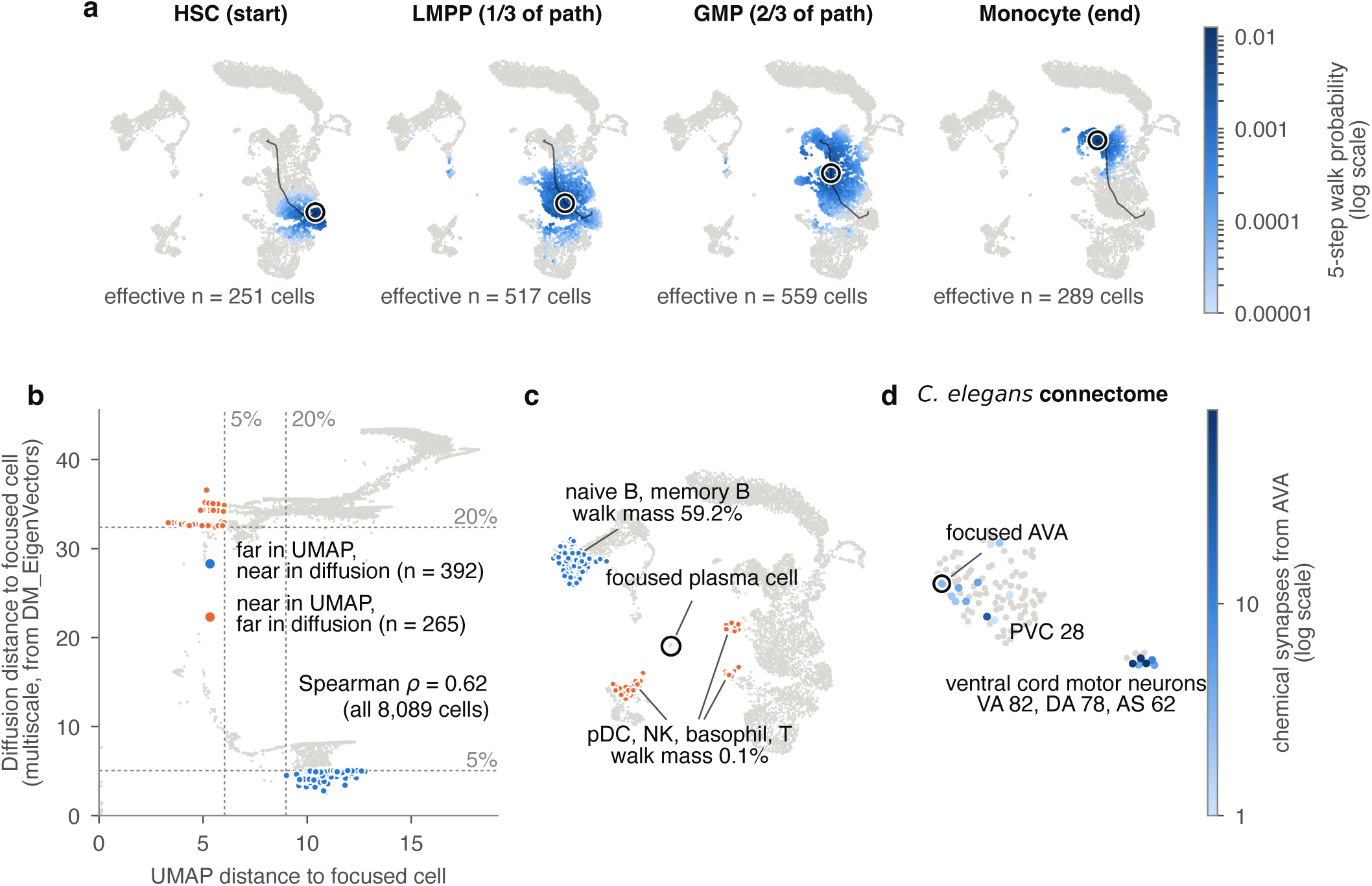
A cell-by-cell matrix, read one focused cell at a time. **a**, The UMAP colored by the focused cell’s row of a dense five-step diffusion walk (`obsp/diffusion_walk_t5`), for four cells along the shortest diffusion path from a hematopoietic stem cell (HSC) to a monocyte (line); the intermediates sit at one and two thirds of the path’s diffusion length. Shared log scale; cells below 10^−5^in gray. *n*_eff_ = 1/ ∑ *p*^2^ is the walk’s effective number of cells. In AnnZarro each panel is one click. **b**, For a focused plasma cell, UMAP distance against multiscale diffusion distance to every other cell (Spearman *ρ* = 0.62). Orange: among its 5% nearest cells on the UMAP but outside its 20% nearest in diffusion space (265 cells); blue: the reverse (392 cells). **c**, The same groups on the UMAP. Cells that look close (pDC, NK, basophil, T) receive 0.09% of the plasma cell’s walk mass; distant-looking naive and memory B cells receive 59.2%. Across all 8,090 cells the UMAP preserves the global rank order of diffusion distances (median per-cell *ρ* = 0.85) but not local neighborhoods (median 14 of 30 nearest neighbors shared); the distortion concentrates in small, isolated populations, while the HSC-to-monocyte trajectory in **a** is faithful (*ρ* = 0.95 for the HSC). **d**, A connectome as a cell-by-cell matrix. *C. elegans* neuron classes on a UMAP of their transcriptomes (CeNGEN ^15^, log TPM), colored by the focused class’s row of chemical synapses (`obsp/chemical_synapses`, synapses sent, summed over the neurons of each class ^14^; log scale), strongest on top; gray, no synapse from the focus. The command interneuron AVA sends its strongest output to the VA, DA and AS motor neurons, which sit in a separate cluster of the UMAP.

**Figure 5:**
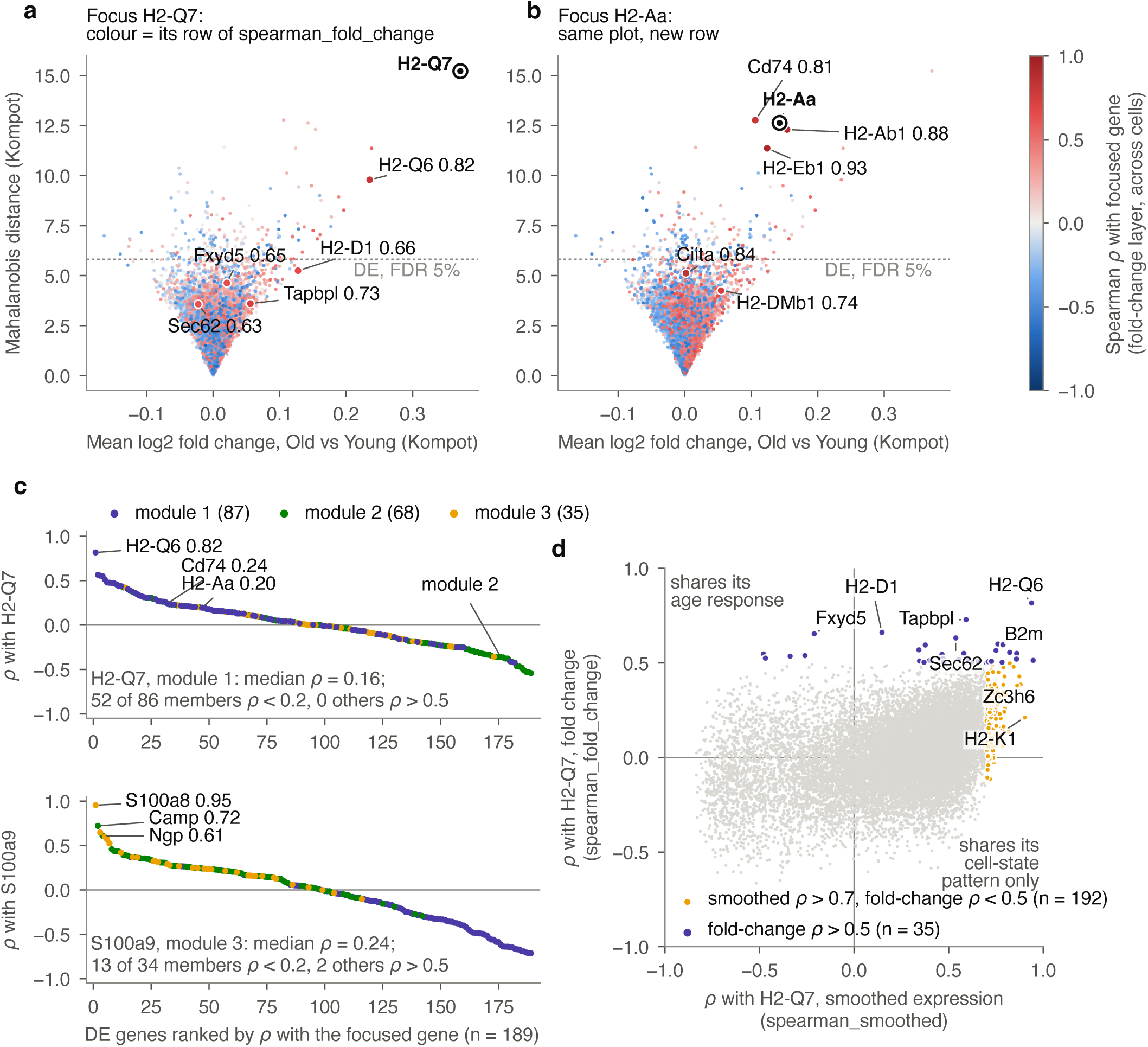
A gene-by-gene matrix over a differential expression volcano. **a**, Kompot volcano (mean log_2_ fold change against Mahalanobis distance, Old vs Young; dashed line, 5% FDR) colored by the focused gene *H2-Q7*’s row of the Spearman correlation of fold changes across cells (`varp/spearman_fold_change`). Its age response is shared by *H2-Q6* (*ρ* = 0.82), *Tapbpl* (0.73), *H2-D1* (0.66) and *B2m* (0.60), MHC class I antigen presentation; only *H2-Q6* is itself called differentially expressed. **b**, The same plot after clicking *H2-Aa* (histocompatibility 2, class II antigen A, alpha): the coloring follows the focus to MHC class II genes (*H2-Eb1* 0.93, *H2-Ab1* 0.88, *Ciita* 0.84, *Cd74* 0.81). **c**, Modules and focused rows. The 190 differentially expressed genes ranked by *ρ* with *H2-Q7* (top) or *S100a9* (S100 calcium binding protein A9; bottom), colored, for comparison, by modules from average-linkage clustering of the same matrix on 1 − *ρ* (*k* = 3, the silhouette maximum; silhouette is flat at 0.23–0.25 for *k* = 2–6). The rows show structure within and across modules: *H2-Q7*’s module holds both MHC class I and class II genes, and 52 of its 86 module-mates have *ρ* < 0.2; *S100a9*’s strongest partners Camp (0.72) and Ngp (0.61) fall outside its module, also at *k* = 5. **d**, *H2-Q7*’s row of the correlation of smoothed expression against its row of the correlation of fold changes. 204 genes share its cell-state pattern (*ρ* > 0.7, smoothed), but only *H2-Q6* also shares its age response; *H2-K1* follows its expression pattern (0.90) but not its change with age (0.21).

**Figure 6:**
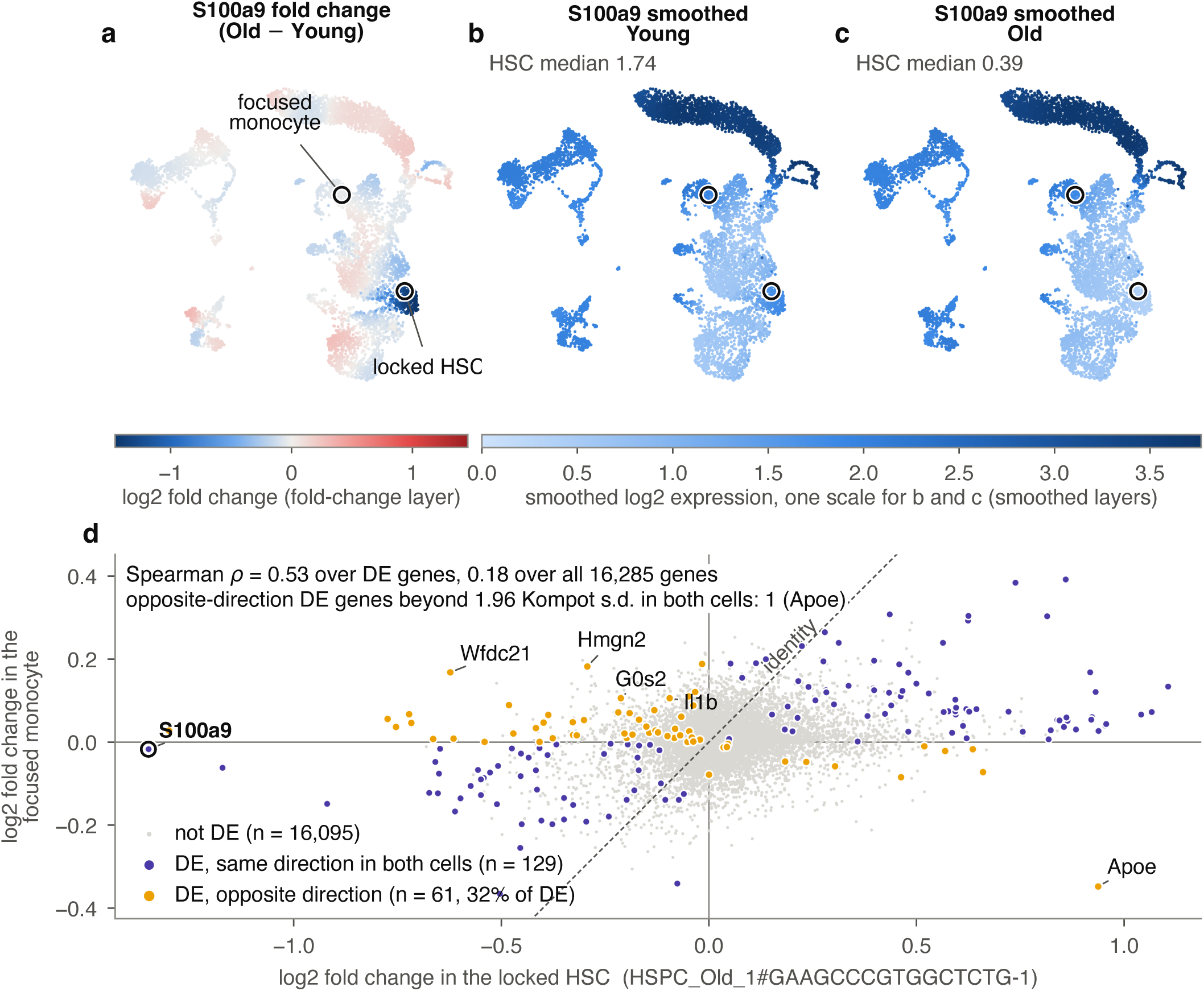
Linking genes to cells and cells to genes. **a**, Focusing *S100a9* colors the UMAP by its column of the Kompot fold-change layer. Its mean log_2_ fold change is −0.05, so it looks unchanged in a volcano, yet it drops in hematopoietic stem cells (median −1.35; next lowest cell type −0.21). **b**, **c**, Its smoothed expression in Young and Old on one shared scale: HSC median 1.74 in Young and 0.39 in Old. **d**, Gene by gene, the fold-change row of a locked HSC against that of the focused monocyte (rings in **a**–**c**). Differentially expressed genes are colored by whether they change in the same (purple, 129) or opposite (amber, 61) direction in the two cells, with no magnitude cut. Opposite-direction changes are mostly small; beyond 1.96 times Kompot’s per-cell standard deviation in both cells, only Apoe changes in opposite directions (up in the HSC, down in the monocyte), against 10 genes changing in the same direction. Spearman *ρ* between the two rows is 0.53 over differentially expressed genes.

**Cell-by-cell relationships.** Any cells × cells matrix in obsp (diffusion or Mellon kernels, kNN connectivities, distances, Kompot posterior covariance, spatial neighbor graphs and distances, or cell-level communication scores) can drive an embedding’s color or axis. Each click on a cell paints that cell’s row over the whole embedding, so walking along a trajectory shows how a kernel or distance function changes with cell state. Palantir pseudotime and branch probabilities ^8^, Mellon densities ^9^ and SEACells metacell assignments ^13^ sit beside these as obs columns. **Fig. 4a** shows such a walk through hematopoiesis. The same view reads a connectome. With synapses between *C. elegans* neuron classes ^14^ stored in obsp and their transcriptomes ^15^ in X, focusing the command interneuron AVA paints its synaptic targets over the transcriptomic UMAP: its strongest targets, the motor neurons of the ventral cord, form a cluster far from it (**Fig. 4d**).

**The geometry of the phenotypic manifold.** Cells occupy a continuous shape in gene-expression space, the phenotypic manifold, and many analyses measure how similar cells are along that shape rather than on a two-dimensional map. Diffusion distances are one such measure: they treat the nearest-neighbor graph as a network, and two cells are close when a random walk passes easily from one to the other. Palantir pseudotime and fate probabilities are computed from them ^8^. Painting the focused cell’s row of such a matrix shows the manifold’s geometry one cell at a time, beside the familiar map: which cells a cell is closest to, and how far its neighborhood reaches. For a plasma cell, most of the walk’s mass lies on naive and memory B cells, its relatives in expression space, although the UMAP draws them far away, and almost none on the pDC, NK, basophil and T cells drawn next to it; along the trajectory from stem cells to monocytes, the map and the manifold agree (**Fig. 4b,c**). An ordering or a fate probability that looks surprising on the UMAP can thus be read in the geometry that produced it.

**Gene-by-gene relationships.** A genes × genes matrix in varp, such as Spearman correlation computed on imputed expression, drives the color of a gene scatter. Coloring a differential expression volcano by correlation to the focused gene shows which genes are co-regulated with it, wherever they sit in effect size and significance, and clicking one of them moves the focus to it. **Fig. 5** shows this for MHC genes in aging bone marrow.

**Differential expression at single-cell resolution.** Kompot ^6^ writes its results across every AnnData slot: per-cell differential abundance in obs; per-gene Mahalanobis distance, mean log fold change and local FDR in var; per-cell-type summaries in varm; dense cells × genes smoothed expression, fold change and fold-change z-score layers; an optional posterior covariance in obsp; and a run history with a field_names map in uns. No single static plot shows these together. In AnnZarro, a gene scatter of Mahalanobis distance against mean log fold change acts as a volcano; clicking a gene recolors linked embeddings with that gene’s per-cell fold change and with smoothed expression in each condition. The reverse direction also holds: clicking a cell recolors the gene scatter with that cell’s fold change for every gene, showing which genes change in that cell state. Locking one cell and refocusing another compares two states gene by gene, which is how opposite regulation of the same genes across cell types becomes visible. **Fig. 6** shows a gene that looks unchanged in the volcano but is lost with age in stem cells.

**Gene sets.** A gene table defines a gene set by any combination of conditions, joined with AND and OR, on the per-gene annotations in var and on columns taken from a chosen gene or cell, such as each gene’s correlation to that gene or its expression in that cell. A set can thus combine thresholds on differential expression statistics, such as one corner of the volcano, with a stored module label, a gene annotation, a name pattern, or similarity and expression thresholds. The optional gene-set panel takes the genes that pass, and the focused gene, to outside services: STRING and g:Profiler for enrichment against the dataset’s genes as background, STRING for the protein network and MyGene.info for what each gene is; Enrichr Reactome and the Human Protein Atlas can be switched on. Changing a threshold and running the panel again gives the next set’s results, and every gene plot linked to the same table shows the set.

While we used the outputs of Palantir and Kompot for illustration, none of these uses needs code specific to Kompot or Palantir. They follow from two general rules: a cell plot can take any per-cell vector, including a row of obsp or a gene column of a layer, and a gene plot can take any per-gene vector, including a row of varp or a cell row of a layer.

### Comparison with other methods

We compared AnnZarro with graphical single-cell viewers, and with Scanpy and Seurat as the baseline of what code can do (**Fig. 1c**). The viewers we compared are CELLxGENE (Explorer and Annotate) ^16,17^, Vitessce ^11^, UCSC Cell Browser ^18^, Single Cell Portal ^19^, Cirrocumulus (part of Cumulus ^20^), SCope ^21^, ShinyCell2^22,23^, Kana ^24^, iSEE ^10^, Loupe Browser, ManiVault Studio ^25^ and VizIt (**Supplementary Note 1**). Viewers built around tissue images, for spatial data and its quality control, serve a different task and were not compared. The paragraphs below follow four of the column groups of **Fig. 1c**: relationships, work, scale and sharing.

**Relationships.** Among these viewers, only ManiVault Studio colors an embedding by a stored cells × cells row. The user selects the row by typing its index and is not shown which cell it belongs to. The other viewers either ignore obsp and varp or reach them only through code. Users cannot color or position points by a genes × genes matrix in any of them, or select a cell or gene in one plot to set the row shown in the others. This link between a focused cell or gene and every panel is the central difference between AnnZarro and these viewers.

**Work.** The tools differ most in what they are for. Many viewers move parts of the analysis into the viewer, so that it can be done without code: CELLxGENE, Cirrocumulus and Loupe edit annotations, and Loupe and CELLxGENE run differential expression on demand. AnnZarro does not try to replace analysis. It is a tool for visualization and exploration: AnnZarro is read-only by design and computes nothing, so every annotation, statistic and matrix it shows is produced in code, where the analysis stays documented and reproducible. Analysis and exploration then form a loop: code computes, AnnZarro shows the results, and what the user sees prompts the next computation. The loop is short because the viewer follows the store. When code, for example in a Jupyter session on the same store, writes new results with anndata’s write_elem, one refresh (Ctrl+R) shows them in the open views, new obs columns, layers and pairwise matrices included, and keeps the layout, the focus and locked colors. On a shared server, the people who run it decide whether stores are updated in place; if they are, every user sees the new results on their next refresh, while the server itself never writes the store. Note that AnnZarro does not render tissue images, as Vitessce does, under the cells, and it cannot export the exact code behind a view, as iSEE does, which would let a user recreate a figure in R.

**Scale.** AnnZarro draws each of the 95.6 million cells of Tahoe-100M as a point (**Fig. 7a,b**). CELLxGENE and Cirrocumulus show datasets of one million or more cells in their documentation and Vitessce millions of points. Since these are demonstrations and do not establish limits, we measured every tool in **Fig. 1c** except CELLxGENE Explorer, Single Cell Portal and VizIt (**Supplementary Note 1**) on datasets built from Tahoe-100M (ManiVault Studio and Loupe Browser by hand, in an interactive desktop session), each drawing a UMAP colored by cell line and by one gene (**Fig. 1c**, Cells strip). The client budget, 16 GiB, covers all browser processes, or the whole process for Scanpy and Seurat; the server budget is a flat 32 GiB for every tool, AnnZarro included. Each tool’s input was prepared in its own format, and preparing it was not measured (**Supplementary Note 1**). AnnZarro was measured on all 95.6 million cells of Tahoe-100M on a laptop (**Fig. 7**): it browses them through a reproducible subset and its parts, and, with the default subset switched off, draws every cell in one plot at the cost of hover and click (benchmarks in **Supplementary Note 2**).

**Figure 7:**
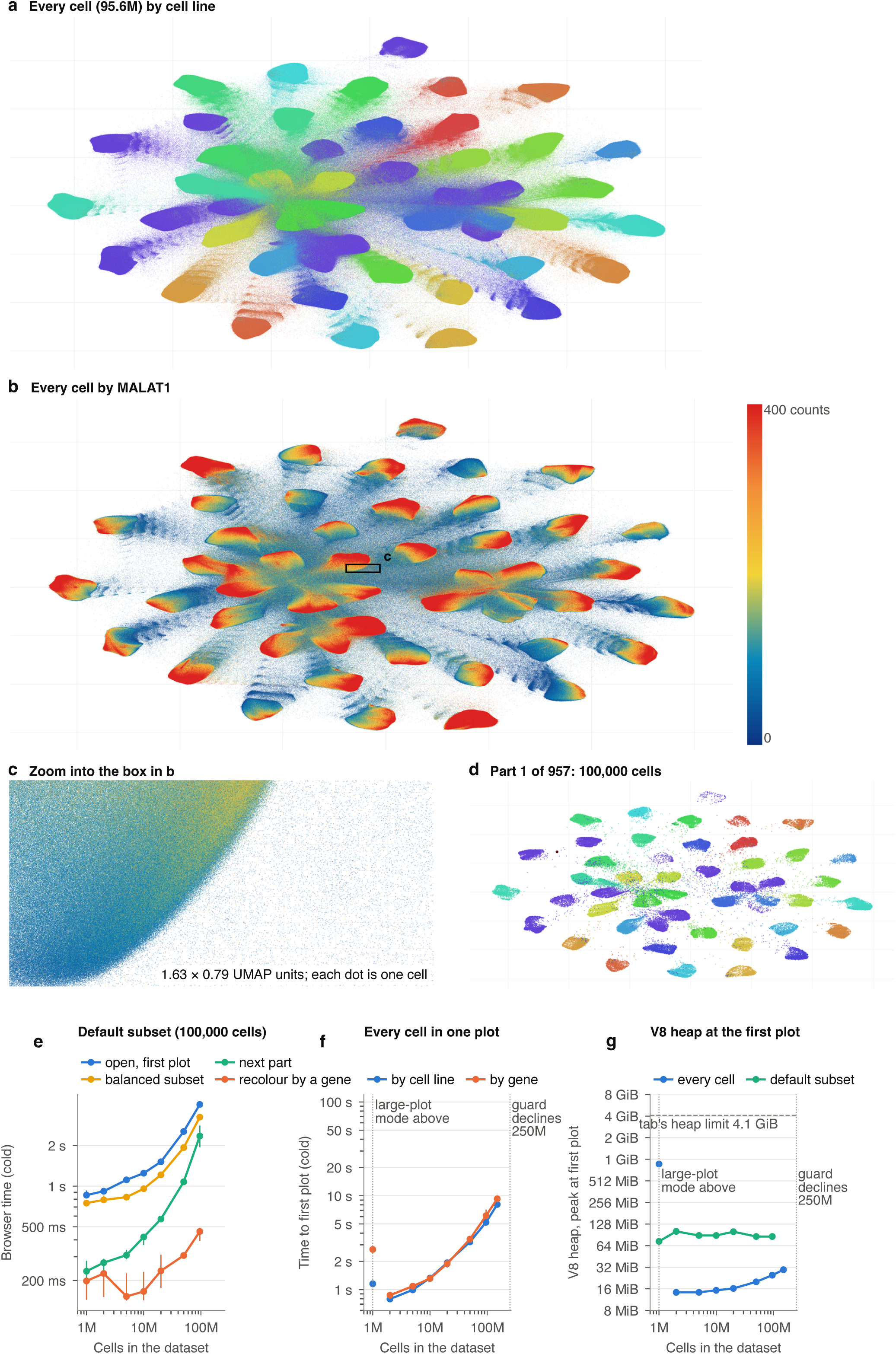
Tahoe-100M on a laptop: every cell in one plot, and a subset that stays interactive. All 95,624,334 cells of Tahoe-100M on a UMAP fitted to 999,073 reference cells and extended to every cell by nearest-neighbor placement (see *Anticipated results*). Headless Chromium 153 on an Apple M3 Max laptop (128 GiB RAM); **a**–**d** show the plot areas of app screenshots. **a**, Every cell colored by cell line in large-plot mode, point size 1 and opacity 0.2; drawn 6.8 s after the page was opened in its screenshot session. In this mode the status line reads “Large plot: no hover/click”, and its tooltip adds that table filters are off. **b**, The same cells colored by *MALAT1* counts, the color scale running from 0 to 400 counts, which 97.9% of cells do not exceed (6.8 s in its screenshot session). Where cells overlap, the highest values are drawn on top. The box marks the region shown in c. **c**, The boxed region of b, 1.63 × 0.79 UMAP units at the edge of one cell line’s island, where 96% of the 574,309 cells belong to that line, opened zoomed from a view link with every cell drawn at the automatic point size and opacity (2.35 px and 0.43): each dot is one cell, colored on the scale of b (drawn in 17.6 s in its screenshot session). The link also focuses one cell; large-plot mode draws no focused-cell marker. **d**, The default subset, part 1 of 957 (100,000 cells), in the regular plot with hover and focus, at the automatic size and opacity, over the same extent as a. **e**, Browser time for actions on the default subset against dataset size, from 1 million cells (prefixes of Tahoe-100M) to all 95.6 million: opening to the first plot, recoloring by a gene, a subset balanced across the 50 cell lines and a step to the next part. Medians over 3 cold runs (2 events per run for recolor and part step), whiskers minimum to maximum; recolor and balanced subset start from a plot colored by a gene. **f**, Time to the first plot with every cell, colored by cell line or by a gene. Datasets up to 95.6 million cells are Tahoe-100M and its prefixes; the larger stores are described in **Supplementary Note 1**. The dotted rule at 1 million points marks the large-plot threshold: the plot at 1 million is regular, larger ones are drawn in large-plot mode; the dotted rule at 250 million marks where the memory guard declines the first plot at default settings, so larger stores are not shown. **g**, Peak JavaScript (V8) heap at the first plot against the tab’s heap limit under default Chromium flags (dashed line). The regular plot fills the heap fastest; large-plot mode keeps the positions off the heap (0.03 GiB at 150 million cells); the default subset stays flat.

**Sharing.** AnnZarro builds on two ideas from other tools. In iSEE ^10^, selecting a row in one table updates the other panels; in AnnZarro, a click on a cell or gene selects its row of every stored pairwise matrix, and each linked panel redraws with it. Vitessce established declarative, URL-shareable view configurations ^11^: a view is saved as a description that reopens it from a link. AnnZarro saves the focused cell and gene in that description too, and users build views by pointing and clicking instead of writing the description by hand.

### Limitations

AnnZarro computes no statistics, clusters, embeddings or correlations. Every quantity it draws from the data must already be stored in the AnnData object, so a gene-module view needs a precomputed varp matrix and a kernel view needs a precomputed obsp matrix.

- **Plot vocabulary.** AnnZarro draws only scatter plots and tables: heatmaps, violins, dot plots and histology image underlays are not available, so users who need them should use scanpy, Seurat or a spatial viewer such as Vitessce alongside. Two plot types, a cell scatter and a gene scatter (2D or 3D), cover all views. Any stored coordinates serve as axes, for cells (UMAP, diffusion components, or spatial positions in obsm, so spatial transcriptomics data are browsed with the same linking) and for genes (any var or varm columns, including gene embeddings).
- **Selection.** Cells cannot be selected by drawing a lasso or box on a plot. Cells are selected instead by table filters on stored values, in arbitrary AND/OR combinations of conditions on any column; one table can hold obs columns, embedding coordinates, gene values and a focused cell’s matrix rows, so any selection fits in one filter. On a large dataset a table filters the cells it has loaded; its conditions on obs columns can also select from every cell, through *Use* in the subset dialog (Step 33). Such a selection is a written rule: it is saved with the panel set and share link, so every user who opens the view gets the same cells, and the rule can be applied to the next sample and reported. A region is then defined by a quantity computed for it, for example each cell’s distance to the nearest tumor cell computed in Python, or in the app a focused cell’s row of a distance matrix; an expert’s annotation of a tissue section enters the same way, as an obs column. On a UMAP or t-SNE, a drawn region would also select cells by positions that distort their similarities ^26^.
- **Storage layout.** A poorly chunked store is slow: latency depends on chunk shape, and a poorly chunked dense layer reads more data per click than needed. AnnZarro is optimized for dense chunked arrays. Sparse matrices are served too, but a gene from a row-compressed (CSR) matrix requires reading all non-zero values, so the Procedure converts X to CSC. Sparse matrices in h5ad files are read the same way, a row or column at a time.
- **Large dense layers.** Each gene click on a dense layer decompresses one chunk column, so chunk shape and float32 storage determine latency. Kompot fold-change and smoothed-expression layers are such dense cells × genes arrays.
- **Very large datasets.** Not every cell of a very large dataset can be explored with hover and click at the same time. Vectors travel as binary float32, sparse rows as their non-zeros and categories as integer codes, but the browser still holds one value per shown cell for each displayed vector. Above 200,000 cells AnnZarro therefore opens on a 100,000-cell subset, and the rest of the cells are reached part by part rather than all at once (Worked example 6). One plot can draw every cell: above 1 million points it switches to large-plot mode, which drops hover, click to focus, table filters, obsp colors and 3D in that plot. Gene plots keep hover and click, and clicking a gene in one still recolors the large plot. The threshold is a server setting; regular plots need far more memory per point, and at its default several of them fit in the browser’s memory side by side. A memory guard checks each plot against the browser’s memory and against the largest plot tested; it allows one plot of every cell up to 200 million cells and declines it at 250 million; with a subset, every store tested opens, up to 1.5 billion cells (**Supplementary Notes 1 and 2**). A billion cells cannot be drawn as interactive points within current browser limits: above the guard’s limit the tab cannot allocate memory for the points’ positions (**Supplementary Note 2**), so datasets of that size are explored on a subset.
- **Data access.** h5ad files must be local, and remote stores need optional dependencies. Local stores and remote zarr stores (`s3: //`, `gs: //`, `https: //`) are served; remote access needs the optional `annzarro[remote]` dependencies and is limited to an allowlist on hosted servers.
- **Reproducibility scope.** A saved view (share link, panel set or exported figure) names its store relative to the data directory and records which cells, genes and fields the store had, so it opens on any server that holds the store (Step 38) and warns when the cells, genes or fields differ. Values changed in place, in the same cells and genes, are not detected. Settings left automatic, such as point size, opacity or an unlocked color range, can differ between AnnZarro versions, and AnnZarro says when a view was saved with another version. With the same store and the same AnnZarro version a view gives the same figure: every exported PNG or SVG carries its view, and `annzarro export --from figure.png` draws it again.
- **Read-only data.** By design, AnnZarro separates producing results, in code, from exploring them: users cannot write labels back to the data or run code on the server; besides small caches of its own, the only state AnnZarro writes is view definitions (panel sets). Curation, as in CELLxGENE Annotate or Loupe, therefore happens in Python, and the store is re-exported.
- **Outside services.** The optional gene-set panel depends on services AnnZarro does not control. Its requests leave from the user’s browser, never from the server, and only after explicit consent, and carry gene identifiers, the species and the dataset’s gene list as background, never cell data; Enrichr and Reactome, which keep what they receive, are off by default.

### Experimental design

The exploration available to users is fixed when the store is written: AnnZarro displays what is stored and computes nothing new. Design therefore happens in four decisions made before the app opens.

#### What to store

Every view reads one AnnData slot (**Fig. 1b**): coordinates and annotations in obsm and obs, expression and Kompot layers in X and layers, per-gene statistics in var and varm, and the two kinds of relationship matrix in obsp (cells × cells: kernels, graphs, distances) and varp (genes × genes: correlations). Category colors come from `uns[‘{key}_colors’]`.

Pairwise matrices are the expensive part. A dense cells × cells matrix grows with the square of cell number (100,000 cells in float32 is 40 GB), so store kNN-sparse kernels or restrict dense obsp matrices to metacells or subsets. A genes × genes matrix over 16,000 genes is about 1 GB in float32; restricting it to highly variable or differentially expressed genes keeps it small.

#### How to store it

Write the store with anndata in zarr format 2, its default. The chunk layout of the dense arrays decides responsiveness (see *Anticipated results*): a gene column of a cells × genes layer costs the chunk’s width in genes times all cells, and a cell row costs the chunk’s height in cells times all genes. Chunks whose aspect ratio follows the matrix’s own, at about 5 × 10^5^ values, keep both fast; anndata 0.12 with zarr 3 already chooses such chunks, and Step 10 rewrites stores that use whole-gene chunks. Pairwise matrices are only read by row, so obsp and varp are chunked in whole rows. Sparse matrices are the secondary case: store them CSC if genes are queried, since CSR forces a read of all non-zero values per gene. For a single machine an h5ad file also opens directly; it is read like a zarr store, but it cannot be opened remotely or rechunked.

#### Where to run it

The same software runs in three arrangements (**Fig. 8**). The desktop app bundles its server for a single user with data on the laptop. A personal server (`annzarro start`) runs where large stores already live, and the browser connects through an SSH tunnel. A lab server hosts datasets for many users behind login and a

**Figure 8:**
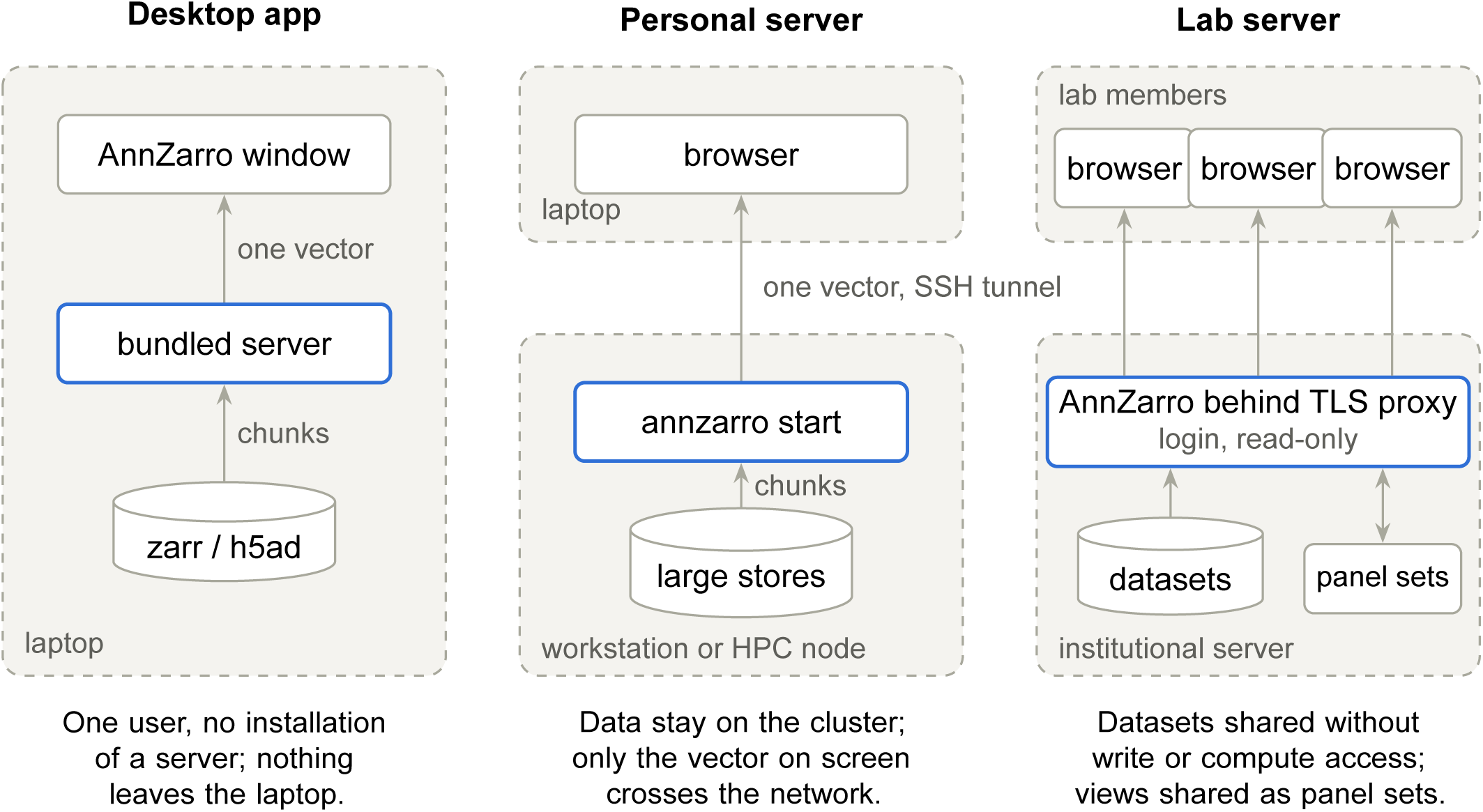
Three ways to run AnnZarro. Dashed boxes are machines. Blue boxes are the AnnZarro server; arrows show what moves per click: compressed chunks from storage to the server, and one decoded vector from the server to each browser.

TLS proxy; it never writes to datasets or runs user code, and panel sets saved by one user are available to all. In every arrangement the browser holds only the vectors on screen and the server holds only the chunks it is reading.

On a lab server, the computational researcher can work on the same stores there, for example in Jupyter. After they write new results with anndata’s write_elem, the refresh button next to the dataset menu shows them, and collaborators see them when they refresh their view (Step 44).

#### Which views to link

Plan panels in pairs that answer one question from both sides: an embedding colored by the focused gene beside a gene scatter colored by the focused cell; or an embedding colored by an obsp row beside a table that masks to a cell state. Lock one focus to keep a reference while moving the other.

#### Expertise needed

Data preparation (Steps 3–10) needs basic Python and familiarity with AnnData. The worked examples (Steps 14–36) and saving and sharing (Steps 37–41) need no programming. Hosting for a lab (Steps 42–44) needs experience administering a Linux server behind a reverse proxy.

### Materials

#### Equipment

- **Client.** Any computer with a current Chromium-based browser or Firefox with WebGL enabled. The browser receives only the vectors on screen and, by default, above 200,000 cells only those of a 100,000-cell subset; a larger subset or every cell can be chosen (Worked example 6). The measurements here used a laptop with 128 GiB RAM (Apple M3 Max); with the default subset the tab’s renderer process used 0.5 GiB at every dataset size up to 95.6 million cells, and with every cell of Tahoe-100M in one plot 8.7 GiB (the default subset switched off, Step 36); the browser’s GPU process added 0.2 GiB and 1.9 GiB. Memory in this protocol is binary (1 GiB = 2^30^ bytes, 1 MiB = 2^20^ bytes), as are installed RAM and memory budgets; data sizes and transfers are decimal (1 GB = 10^9^ bytes, 1 MB = 10^6^ bytes), and quoted app messages keep their own units. Process memory is resident memory (RSS), except in the tool comparison (**Supplementary Note 1**), which reports physical footprint, a different measure.
- **Server (optional).** A Linux workstation or HPC node with read access to the zarr stores, Python 3.9 or later and an open port reachable from the client, directly or via an SSH tunnel. Memory scales with the largest chunk read, not the dataset size. Serving the default subset of Tahoe-100M (95.6 million cells) the server process used at most 1.7 GiB in the browser session and 2.6 GiB under eight concurrent subset requests; with the subset switched off, drawing and recoloring every cell in one plot took up to 7.0 GiB.
- **Desktop alternative.** The standalone app for macOS, Windows or Linux bundles the server and runs entirely on one machine with no separate installation.

#### Software

**Table 1.** Software used in the Procedure. Minimum versions, the versions tested and where to obtain each package.

| Package | Version | Role | Source |
| --- | --- | --- | --- |
| AnnZarro | 0.4.1 (tag v0.4.1, commit 78c9da6) | Server, web client, CLI | PyPI (pip install annzarro), GitHub, standalone apps on GitHub releases |
| anndata | 0.10 or later (tested 0.12.19) | Writing the zarr store | PyPI |
| zarr-python | 2.13 or later (tested 3.1.6) | Storage | PyPI |
| scanpy | 1.10 or later (tested 1.11.5) | Preprocessing, embeddings, neighbor graphs (optional) | PyPI |
| kompot | 0.8 or later (tested 0.8.0) | Differential abundance and expression (optional, for the case study) | PyPI, bioconda |
| Palantir | 1.4 or later (tested 1.4.5) | Diffusion maps, pseudotime, fate probabilities (optional) | PyPI |
| Mellon | installed with kompot (tested 1.7.1) | Cell-state densities (optional) | PyPI |

AnnZarro’s runtime dependencies (Flask, Flask-CORS, Werkzeug, zarr, numpy, scipy, h5py, psutil, PyYAML) install automatically. The Procedure was tested with Python 3.11 and the versions given.

#### Example data

Demonstrations use the processed dataset of the Kompot tutorial: CITE-seq of murine bone marrow across aging (Young, Mid, Old), deposited as murine_bone_marrow_aging.h5ad on Zenodo (2.4 GB, CC-BY 4.0). The Kompot getting-started notebook turns it into bm_aging_processed.h5ad (Step 4), which carries differential abundance in obs, gene statistics in var, and fold-change and smoothed-expression layers. Steps 3–10 convert this file to an AnnZarro-ready zarr store and add a gene-gene Spearman correlation in varp and a diffusion kernel in obsp.

### Procedure

**Installation (Steps 1–2)**

**TIMING** 5–10 min

1. Install AnnZarro on the machine that will read the data. Follow option A to install the Python package, which also runs data preparation (Steps 3–10), or option B to install the desktop app, which views prepared stores without a Python installation.

A. Python package

i. Create a fresh environment and install AnnZarro with the packages used for data preparation: 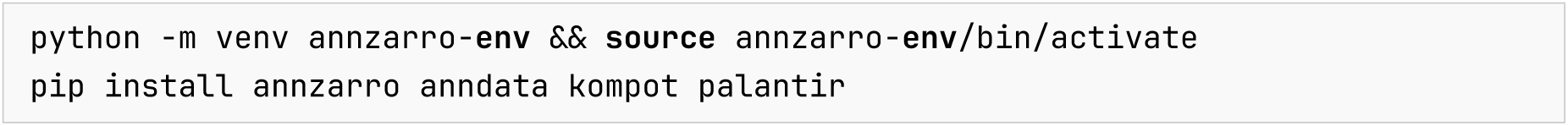
B. Desktop app

i. Download the app for macOS, Windows or Linux from the AnnZarro GitHub release page and install it as usual for the operating system. It bundles the server. Data preparation (Steps 3–10) still needs the Python packages of option A, on this or another machine.
2. Check the installation. For option A, run `annzarro --help`; the subcommands start, stop, user, install, config, desktop and export should be listed. For option B, open the app; the data directory is chosen in it at Step 11.

**Data preparation (Steps 3–10)**

**TIMING** about 10 min on a 16-core workstation; 32 GiB RAM

3. Download the murine bone marrow aging data (2.4 GB) from Zenodo ^27^ and verify its checksum. The script docs/_tools/datasets/bm_aging.py in the AnnZarro repository (v0.4.1) does both and, run without options, also performs Steps 4–10; it pins the tested package versions (Software). 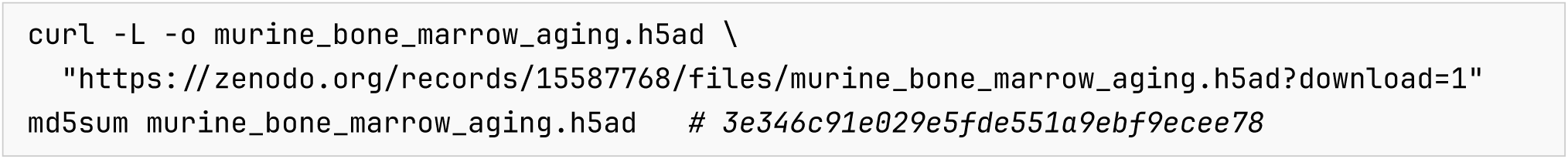
4. Run Kompot differential abundance and differential expression between Young and Old, following the Kompot getting-started tutorial (the `kompot` stage of `bm_aging.py`; about 1 min). Write the result to `bm_aging_processed.h5ad`. **CRITICAL STEP** Do not call kompot.cleanup(adata) before saving, as the tutorial does: it deletes the smoothed-expression and fold-change layers that Step 7 and Worked examples 3 and 4 need. **PAUSE POINT** The processed h5ad (5.6 GB) can be stored indefinitely.
5. Load the processed file and read the keys Kompot wrote from its run record. Use these names in later steps rather than typing them; they differ between Kompot versions. 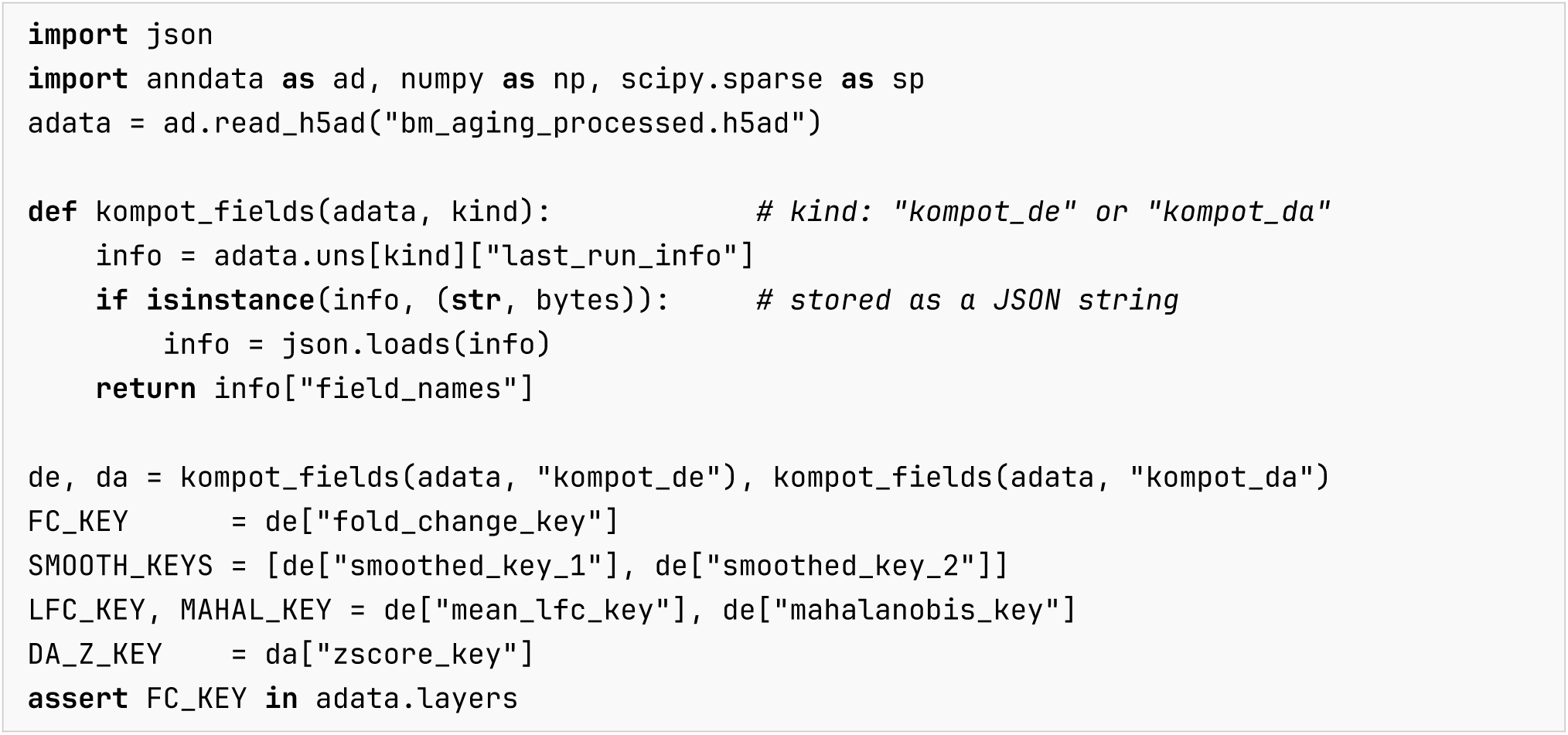 **CRITICAL STEP** `field_names` also lists keys that were not stored (for example a posterior covariance when it was not requested). Check membership before using a key. **? TROUBLESHOOTING** (Table 2)
6. Add a dense cell-by-cell similarity. Palantir’s diffusion maps already store a sparse kernel (`DM_Kernel`) and its row-normalized transition matrix (`DM_Similarity`) in obsp, but one row of a kNN kernel covers only about 33 cells and appears as a speck on the embedding. A few steps of the diffusion random walk spread each row over a few hundred cells and make the cell-state neighborhood visible: 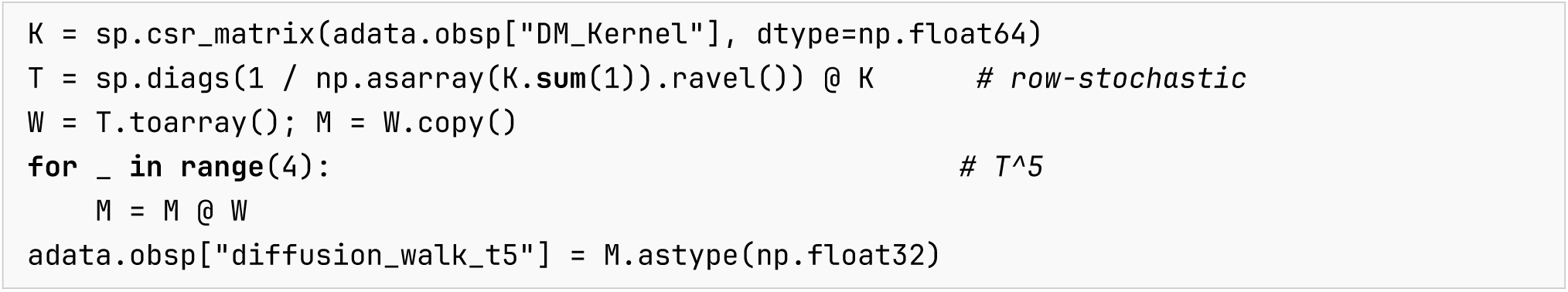 **CRITICAL STEP** A dense cells × cells matrix costs 4*n*^2^ bytes (0.26 GB for 8,090 cells, 4 GB for 32,000). Above roughly 30,000 cells, keep the sparse kernel or threshold *T^t^* into a sparse matrix; AnnZarro reads sparse rows lazily.
7. Add gene-by-gene Spearman correlations. Correlation of the fold-change layer asks which genes’ age effects vary together across cell states and recovers co-regulated aging responses; correlation of smoothed expression recovers cell-type programs. Store both (about 30 s and 4 GiB of RAM for 16,285 genes): 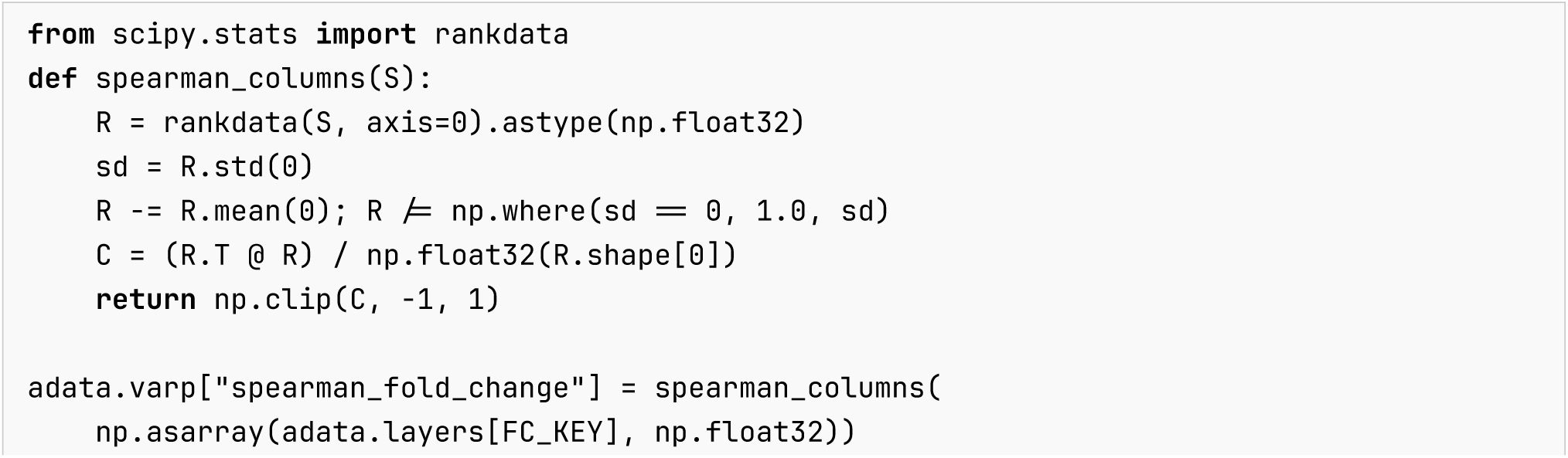
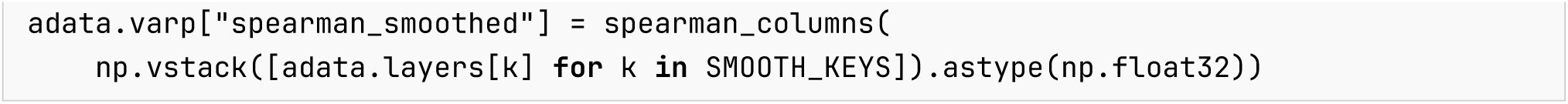 **CRITICAL STEP** A genes × genes float32 matrix costs 4*n*^2^_genes_ bytes (about 1 GB each here). Restrict to highly variable or differentially expressed genes if memory is short.
8. Cast dense arrays to float32, which halves storage and the data decompressed per click, and store sparse matrices that will be read by gene in compressed sparse column (CSC) form. From CSC a single gene is one contiguous slice; from the default compressed sparse row form AnnZarro has to scan every stored index to find one gene’s values, which is far slower on large data. 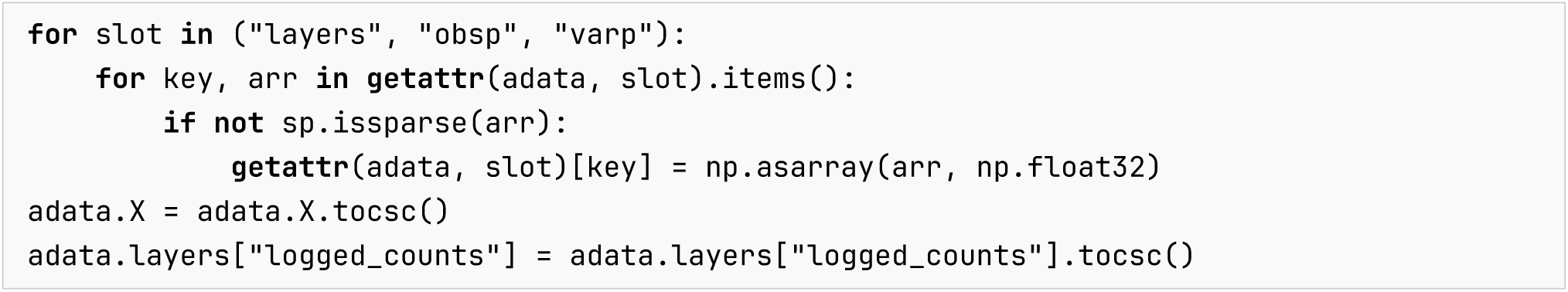 In this dataset X holds scaled expression; color cells by logged_counts for interpretable expression values.
9. Write the store in zarr format 2, anndata’s current default, which every AnnZarro server reads. 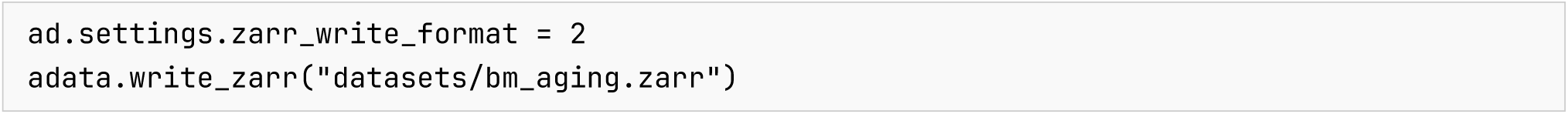
10. (Optional) Check chunk shapes of large dense arrays. anndata 0.12 with zarr 3 already chooses suitable chunks; rewrite only stores produced elsewhere with whole-gene or square chunks. Showing a gene across cells reads one column of a cells × genes layer, and showing a cell across genes reads one row; the cost of each grows with the chunk’s extent along the other axis. Chunks whose aspect ratio follows the matrix’s own (*n*_cells_/*n*_genes_) and hold about 5 × 10^5^ values keep both reads fast at any size. Pairwise matrices in obsp and varp are only read by row, so their chunks span whole rows. Because `write_zarr` consolidates the store’s metadata, open it without consolidation, rewrite, and re-consolidate: 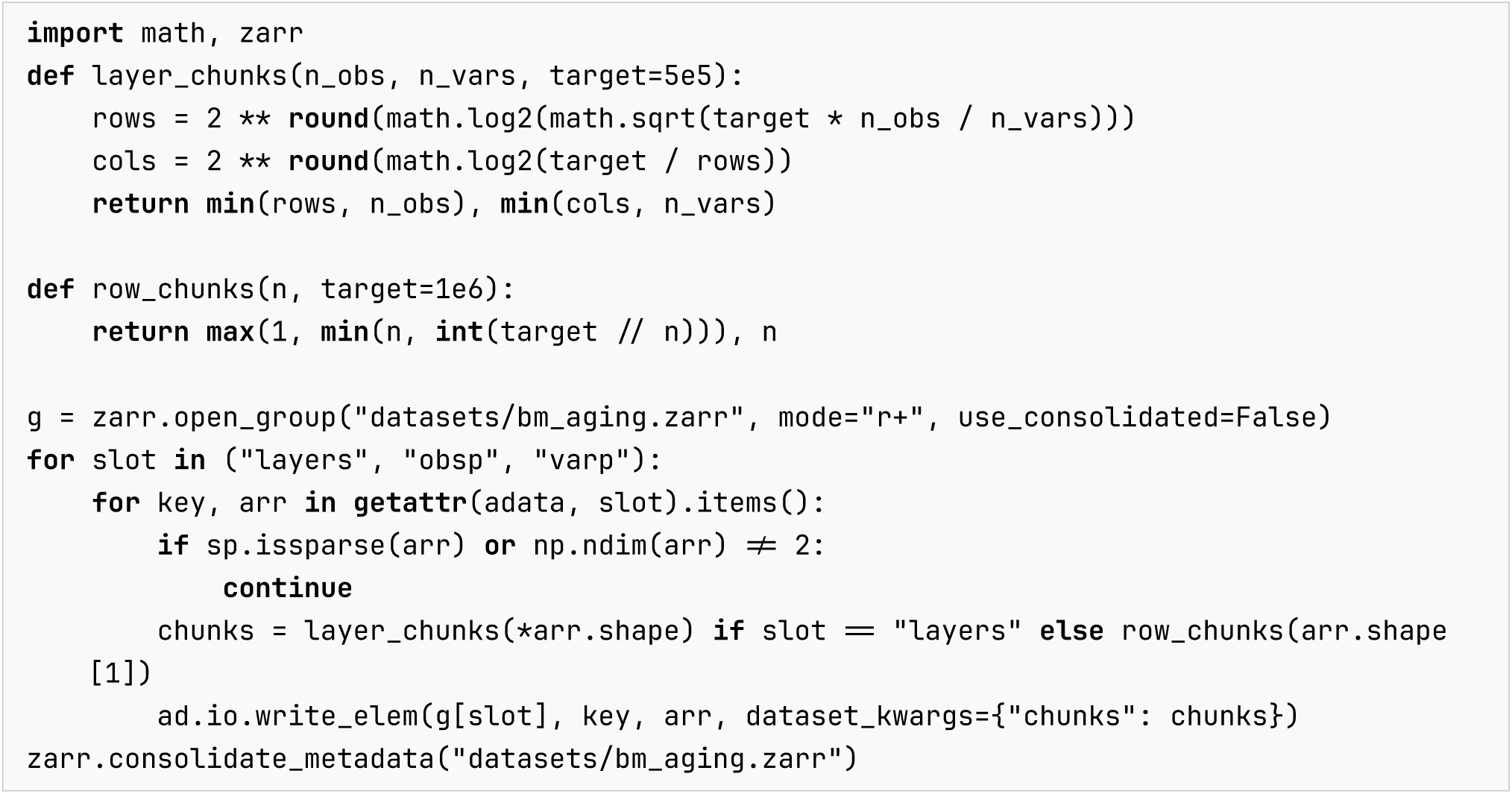 For the demonstration data (8,090 cells, 16,285 genes) this gives 512 × 1024 chunks for layers; for 1 million cells and 5,000 genes it gives 8192 × 64. Whole-gene chunks such as (*n*_cells_, 4), common in practice, make a single cell row decompress the entire layer: 7.8 s for 1 million cells, against 0.38 s with the rule’s 8192 × 64 chunks (**Supplementary Note 2**). **CRITICAL STEP** Skipping consolidate_metadata leaves a store that neither AnnZarro nor anndata can read. **? TROUBLESHOOTING** (Table 2)

**Starting AnnZarro (Steps 11–13)**

**TIMING** 2 min

11. Start the server on the directory that holds the store. 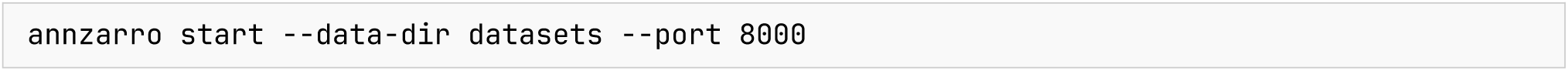 On a laptop the browser opens at http://localhost:8000 without a login. On a remote host, either open an SSH tunnel (`ssh -L 8000:localhost:8000 host`) or run a multi-user server (Steps 42–44). In the desktop app, choose the data directory in the app instead.
12. Select `bm_aging.zarr` in the *Dataset* menu. The status bar reports 8,090 cells and 16,285 genes; the header shows the *Focused Gene* and *Focused Cell* selectors with back and forward buttons. **? TROUBLESHOOTING** (Table 2)
13. Add panels with *Create New Panel* (*Cell Plot*, *Gene Plot*, *Cell Table*, *Gene Table*). Drag the dividers to arrange them; *Duplicate* or *Reopen Panel* clones a configured panel. Ready-made panel sets and share links for every worked example are on the documentation’s procedure page (https://annzarro.readthedocs.io/en/latest/paper/procedure.html#paper-procedure-examples). In *Load Panel Set*, *Load* opens the set’s dataset and every panel that was open when it was saved, in its layout; the icons beside it load the set on the open dataset, or only add its panels to the closed list.

**Worked example 1: walk a diffusion neighborhood across the embedding (Steps 14–17)**

**TIMING** 5 min

**Tutorial:** https://annzarro.readthedocs.io/en/latest/tutorials/cell-similarity.html

14. Create a *Cell Plot*. Set *X* and *Y* to *obsm* → `X_umap`, columns 0 and 1. Set *Color* to *obsp* → `diffusion_ walk_t5`. The row selector reads *Focused cell*.
15. Click a hematopoietic stem cell (for example `HSPC_Old_1#GAAGCCCGTGGCTCTG-1`). It becomes the focused cell everywhere, and the plot recolors by that cell’s row: the cells its diffusion walk reaches in five steps. Only this one row is fetched; the embedding is not.
16. Click cells along the trajectory toward monocytes and watch the neighborhood move and change shape. Use the history arrows beside *Focused Cell* to step back.
17. (Optional) Split the view, pick this panel under *Duplicate* or *Reopen Panel*, and color the copy by `DM_ Kernel` or `distances` to compare neighborhood definitions for the same cell. For spatial data, set *X* and *Y* to the spatial coordinates in obsm.

**Worked example 2: browse co-regulated genes in the differential expression volcano (Steps 18–21)**

**TIMING** 5 min

**Tutorial:** https://annzarro.readthedocs.io/en/latest/tutorials/gene-similarity.html

18. Create a *Gene Plot*. Set *X* to *var* → `kompot_de_Young_to_Old_mean_lfc` and *Y* to *var* → `kompot_ de_Young_to_Old_mahalanobis`. This is the volcano. **? TROUBLESHOOTING** (Table 2)
19. Set *Color* to *varp* → `spearman_fold_change`. The row selector reads *Focused gene*.
20. Click a strongly changed gene, for example *H2-Q7*. It becomes the focused gene, and every gene is colored by how similarly its age effect varies across cell states. Genes of the same response light up wherever they sit in effect size; for *H2-Q7* these include *H2-Q6*, *Tapbpl*, *H2-D1* and *B2m* (MHC class I antigen presentation). **CRITICAL STEP** Start from a gene that is in the correlation matrix. If the focused gene is absent, every point is gray. **? TROUBLESHOOTING** (Table 2)
21. Click a correlated gene to move the focus to it, or switch *Color* to `spearman_smoothed` to see which genes share the focused gene’s cell-type expression pattern instead. To fade weak correlations, set *Map* to Blues, press *Reverse* so that the pale end is low, enter 0.3 as *Min* and press *Lock Range*: genes below 0.3, anti-correlated ones included, take the pale color, and the range holds as the focus moves.

**Worked example 3: from a gene to the cells where it changes (Steps 22–24)**

**TIMING** 5 min

**Tutorial:** https://annzarro.readthedocs.io/en/latest/tutorials/cells-and-genes.html

22. Add a *Cell Plot* on `X_umap`. Set *Color* to *layer* → the Kompot fold-change layer, column *Focused gene*. Enable *Center at 0* and a diverging color scale. **? TROUBLESHOOTING** (Table 2)
23. Add two more *Cell Plot* panels colored by the smoothed expression layers of Young and Old, and lock their color ranges to a shared range. **? TROUBLESHOOTING** (Table 2)
24. Click genes in the volcano of Worked example 2. All three embeddings follow, showing where in state space each gene changes and in which direction.

**Worked example 4: compare two cell states gene by gene (Steps 25–27)**

**TIMING** 5 min

**Tutorial:** https://annzarro.readthedocs.io/en/latest/tutorials/cells-and-genes.html

25. Create a *Gene Plot*. Set *X* to *layer* → the fold-change layer, row *Focused cell*, and press the lock button: *X* stays on the current cell (for example a stem cell). **? TROUBLESHOOTING** (Table 2)
26. Set *Y* to the same layer, row *Focused cell*, unlocked. Set *Color* to *varp* → `spearman_fold_change`, row *Focused gene*.
27. Click a cell of another type in any embedding. Genes on the diagonal change alike in both states; genes off it change differently, and genes in the upper-left or lower-right quadrants change in opposite directions.

**Worked example 5: define subsets with tables and mask plots (Steps 28–31)**

**TIMING** 5 min

**Tutorial:** https://annzarro.readthedocs.io/en/latest/tutorials/tour.html

28. Create a *Cell Table*. In the column picker, tick the cell type column and the Kompot differential abundance z-score in obs, then click *Apply Changes*.
29. Filter with the search box (for a regular expression, press the *.\** button under it and type, for example, `Mid|Old`), or with *Advanced Search*, which combines conditions with AND and OR, for example cell type equals HSC AND abundance z-score above 2, the filter of **Fig. 2c**.
30. In any plot’s options, set *Table* to this table. Cells outside the filter turn gray; the eye toggle hides them instead. Masking happens in the browser, with no new data requests.
31. Export the filtered rows as CSV. Click a cell ID in the table to focus that cell.

**Worked example 6: datasets of millions of cells (Steps 32–36)**

**TIMING** 5 min

**Tutorial:** https://annzarro.readthedocs.io/en/latest/tutorials/millions.html

Steps 32–36 apply to any dataset of more than 200,000 cells. The times quoted are for Tahoe-100M (95,624,334 cells ^7^) on a laptop (**Fig. 7**). Worked examples 1–5 work unchanged on a subset.

32. Open the dataset. It opens on a subset of 100,000 cells with seed 0, and the statistics bar shows the badge *Subset* ⋅ *seed 0*. Every panel and table shows the same cells. The selection is fixed by the seed and each cell’s row, so the same seed gives the same cells on every machine, and a larger subset with the same seed contains the smaller one.
33. (Optional) Click the badge to change the subset: the number of cells, the seed, *Balanced across* an obs column (for example the cell line, which gives small groups an equal share) and *Filter*: conditions on obs columns joined by AND (*Add condition*), or a cell table’s filter (*Use*), which brings its AND/OR groups and text conditions such as contains or starts with. The preview line states how many cells pass and how many are taken per group. Click *Apply*; every panel reloads. On Tahoe-100M, opening the dialog, choosing *Balanced across* the cell line and applying it takes 3.3 s.
34. Step through the rest of the cells with the part buttons next to the badge, or type a part number and press Enter. The parts are disjoint and together hold every cell that passes the conditions once; Tahoe-100M in parts of 100,000 has 957. Layout, colors, zoom and locks stay as they are. Each step takes 2.8 s on Tahoe-100M. A line under each plot counts the cells it shows and says why the others are not drawn, for example that they are not in this part or are hidden by a table filter (cells a filter only grays out still count as shown).
35. To focus a cell outside the current part, type its name into the *Focused Cell* picker; cells outside the subset follow the subset’s cells, tagged not shown. A focused or locked cell stays focused when users step to another part: its rows of obsp matrices and layers are still read and painted over the part’s cells, and the header marks it as not in this part. Share links and panel sets record the part and the focus, and reopen both. **? TROUBLESHOOTING** (Table 2)
36. (Optional) To draw every cell, switch off *Show a subset of the cells* in the badge dialog. Above 1 million points a Cell Plot switches to large-plot mode, and the status line under the plot reads *Large plot: no hover/click*. Hover, click to focus, the focused-cell marker, table filters, obsp axes and colors and 3D are off; pan, zoom, point size, opacity and color by an obs column or a gene remain. Until set by hand, point size and opacity follow the number of points in view, so zooming in enlarges the points. On Tahoe-100M every cell is drawn in 5–7 s at the automatic size and opacity, and in 5–6 s at point size 1 and opacity 0.2, which shows density instead of a solid shape (**Fig. 7a,b**). Switch the subset back on to recover hover and focus. **? TROUBLESHOOTING** (Table 2)

**Saving, sharing and exporting (Steps 37–41)**

**TIMING** 5 min

37. Click *Save Panel Set* and name it. The set is stored on the server and appears under *Load Panel Set* for every user of this instance. **? TROUBLESHOOTING** (Table 2)
38. To move a set to another server, click its download icon in the *Load Panel Set* dialog to save it as JSON, and add it there with *Upload file*. It opens on any server whose data directory holds the same store; the badge on its card says whether that store was found.
39. Click *Share Link* in the header to copy a link that holds the dataset, the layout with every panel’s settings, the cell subset and the focused cell and gene. The view is stored in the link’s fragment, which the browser never sends to the server, so the server sets no limit on its length.
40. For figures, open *Plot Options* in a panel, set fonts, axes and size, and export SVG, PNG, JPEG or WebP.
41. Record the panel set name, the dataset version and the AnnZarro version in the figure legend or methods. Exported PNG and SVG files also carry the view they show, the store’s identity and the AnnZarro version.

**(Optional) Hosting for a lab (Steps 42–44)**

**TIMING** 30 min

42. On a shared server, create accounts with `annzarro user add --username NAME`; add `--admin` for the person who manages the server (`annzarro user set-admin` changes an existing account).
43. Start the server with `annzarro start --data-dir/shared/datasets --host 0.0.0.0 --port 8000 -- detach`; login is enforced for non-local hosts. Run it behind the institution’s reverse proxy with TLS. **? TROUBLESHOOTING** (Table 2)
44. Give the server account read-only access to the data directory. AnnZarro never writes to datasets and runs no user code; it writes only panel sets and small caches of its own, outside the data directory. Admins can also open datasets elsewhere on the server; to keep everyone inside the data directory, set `server.arbitrary_paths: local-only` in the configuration. When a dataset changes on disk, users see the new values with the refresh button next to the dataset menu (Ctrl+R); the browser’s reload can keep showing old values.

### Troubleshooting

**Table 2.**
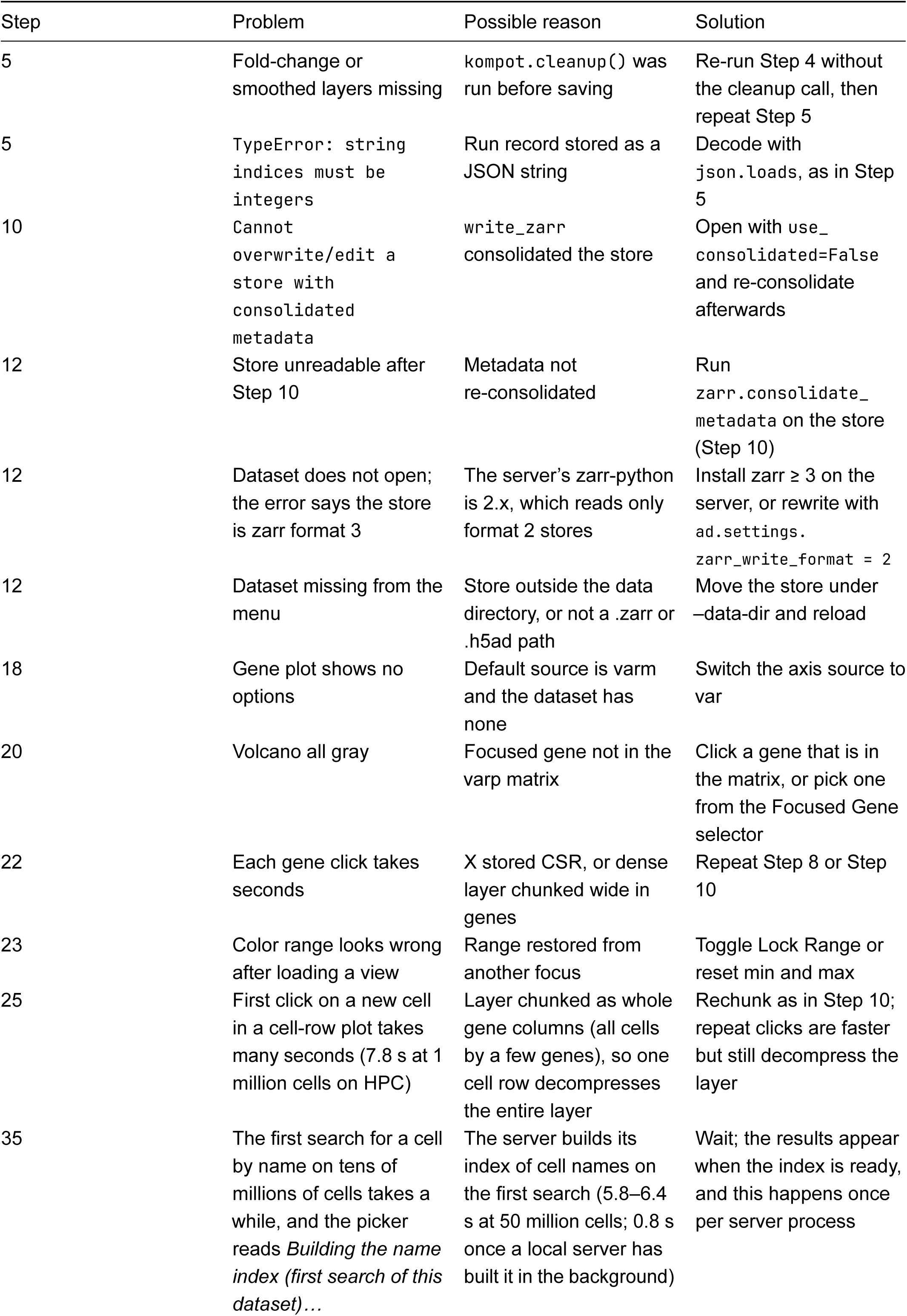
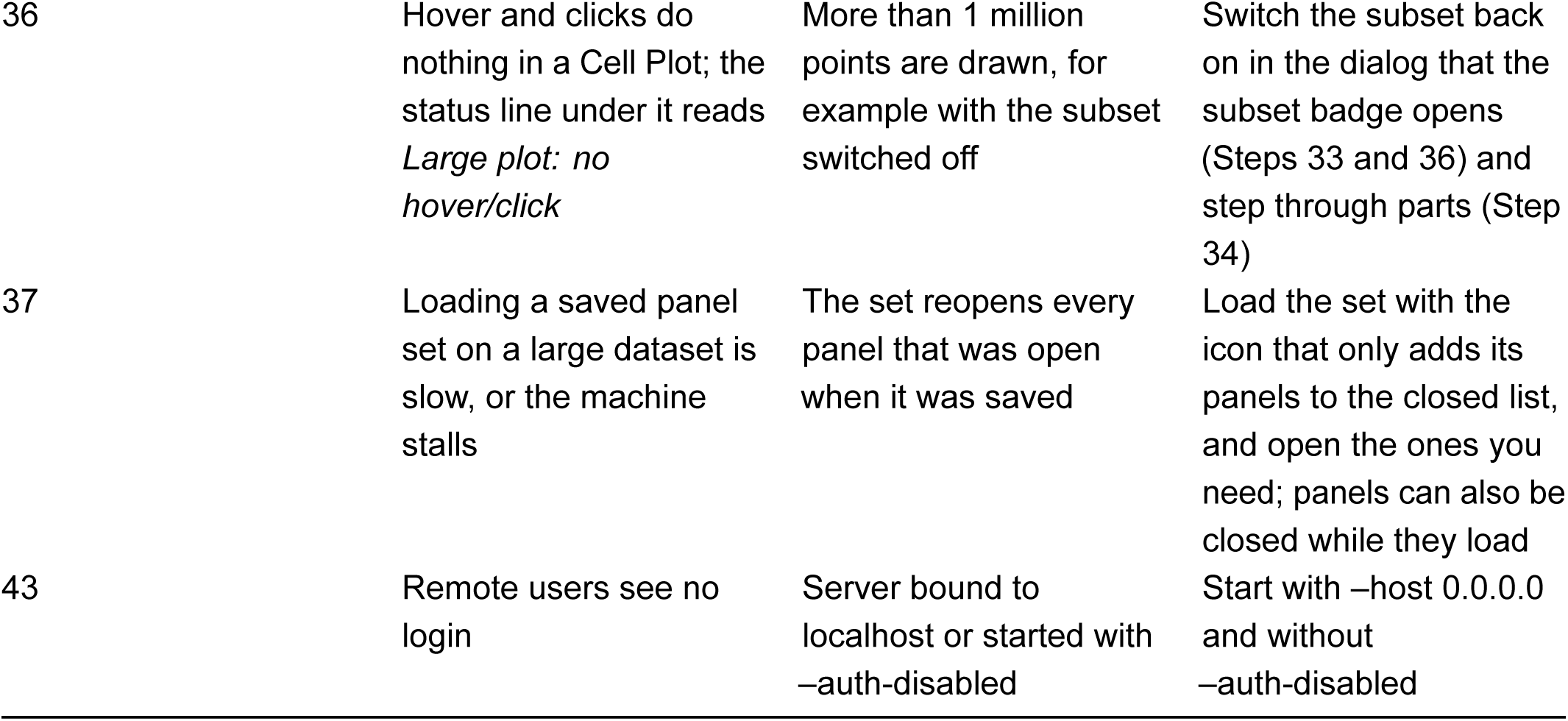
Troubleshooting. The step where each problem is first observed, its possible reason and the solution.

### Timing

Data preparation was timed; the other times are the authors’ estimates for a first run.

- Steps 1–2, installation: 5–10 min
- Steps 3–10, data preparation: about 10 min on a 16-core workstation with 32 GiB RAM (download 5.5 min, Kompot 1 min, store preparation 1 min)
- Steps 11–13, starting AnnZarro: 2 min
- Steps 14–31, Worked examples 1–5: about 5 min each to set up; exploration itself is open-ended
- Steps 32–36, Worked example 6, datasets of millions of cells: 5 min
- Steps 37–41, saving and export: 5 min
- Steps 42–44, lab hosting: 30 min

### Anticipated results

What a user sees first: running Worked examples 1–5 on the bone marrow data reproduces the views in **Figs. 4–6**:

- **Diffusion walk (Worked example 1, Fig. 4a).** A stem cell’s five-step walk stays within its cell type (91% of its mass); moving the focus to a lymphoid-primed or granulocyte-monocyte progenitor spreads it across 517–559 effective cells and over two or more cell types, and at the monocyte it contracts again.
- **Neighbors on the manifold (Fig. 4b,c).** Coloring by a plasma cell’s diffusion row shows most of its walk on naive and memory B cells that look distant on the UMAP, and almost none on the nearby-looking pDC, NK, basophil and T cells; for the stem cell, UMAP and diffusion distances agree. The tutorial linked in **Fig. 4** reproduces both panels in the app.
- **Co-regulated genes in the volcano (Worked example 2, Fig. 5).** Focusing *H2-Q7* highlights the MHC class I antigen-presentation response (*H2-Q6*, *Tapbpl*, *H2-D1*, *B2m*); clicking *H2-Aa* switches the highlight to class II genes. Most of these partners are not themselves called differentially expressed. For comparison, **Fig. 5c** groups the same correlations into modules by hierarchical clustering. The modules give a coarse overview, with no number of modules standing out, and the focused rows add the

detail within them: *H2-Q7*’s module holds both class I and class II genes, which its row separates, and *S100a9*’s strongest partners sit in another module. Switching the color to the correlation of smoothed expression (Step 21) asks a different question: many genes share *H2-Q7*’s expression pattern across cell states, but few share its change with age (**Fig. 5d**).

- **Gene-to-cell maps (Worked example 3, Fig. 6a–c).** *S100a9*, unchanged on average, is lost with age in stem cells only.
- **State comparison (Worked example 4, Fig. 6d).** A locked stem cell has much larger age effects than a monocyte (median | log_2_ FC| over differentially expressed genes 0.41 against 0.068); the two agree in direction for 129 of 190 differentially expressed genes.
- **Masking (Worked example 5).** A table filter on cell type and abundance z-score grays out all other cells in every linked plot without new requests.

A saved panel set reproduces the same five views for any other user of the instance, and SVG export provides figure-ready versions.

**Good and problematic results.** Each click transfers one vector, never a matrix (**Fig. 9**), so the cost of an interaction grows with the number of cells or genes and not with the size of the stored matrix. Three signs tell a good result from a problematic one.

- **Good: a click redraws at once.** In a store written as in Step 10, focused rows and columns arrive in well under a second; on the demonstration store a cell row of the fold-change layer is read from the store in 10 ms.
- **Problematic: a click on a cell takes seconds.** The layer is then chunked as whole gene columns, so one cell row decompresses the entire layer. In the HPC benchmark, a cells × genes layer of Tahoe-100M (1 million cells, 5,000 genes) in chunks of all cells by 64 genes took 8.7 s cold for one cell row, against 0.38 s in the Step 10 shape for this layer, 8192 × 64 (**Fig. 9d**). The status line under the plot shows the wait; rechunk as in Step 10 (Troubleshooting, Step 25).
- **Problematic: every point is gray in a plot colored by a matrix row.** The focused cell or gene is not in that matrix, for example a gene outside the correlated gene set; focus one that is (Troubleshooting, Step 20).

**Figure 9:**
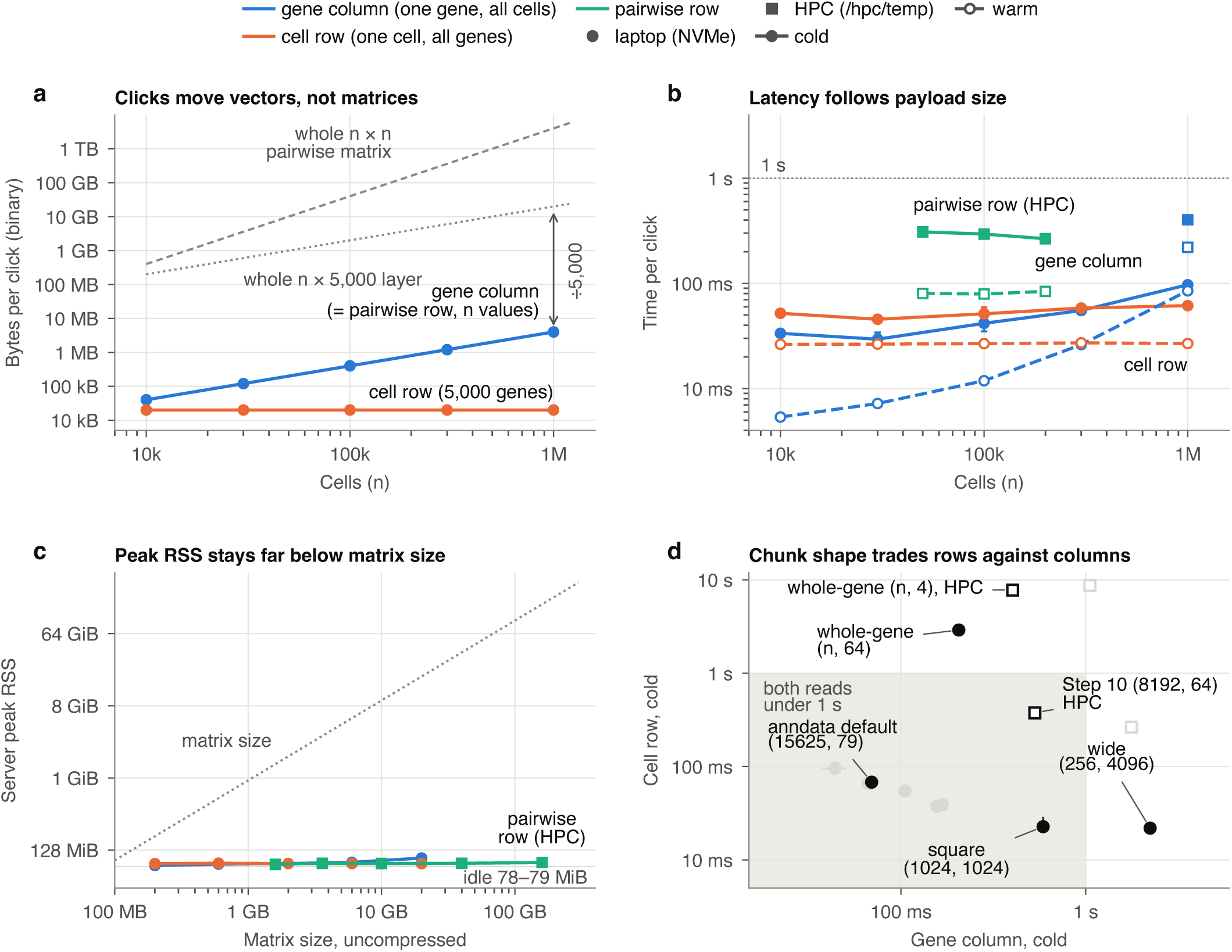
Interactions cost one vector, and memory stays flat. **a**, Bytes sent per click against the number of cells: a gene column, a cell row (5,000 genes) and a row of a dense pairwise matrix, with the size of the whole layer and the whole pairwise matrix for reference. Bytes are computed from the binary format, 4 bytes per float32 value, for the slices read in b (every value of these benchmark arrays is non-zero, so none is sent sparse). At one million cells a 20 GB layer is shown through a 4 MB gene column, one part in 5,000. **b**, Latency per click on a laptop (M3 Max, NVMe), cold (filled) and warm (open; a new index). Gene columns stay under 1 s to one million cells on the laptop. On HPC nodes (Xeon Gold 6254, a whole 36-CPU node; data on /hpc/temp, an NFSv4.1 scratch filer), a gene column of a dense Tahoe-100M layer (whole-gene chunks) takes 0.40 s cold at 1 million cells, and a row of a dense 200,000 × 200,000 matrix (160 GB uncompressed) 0.27 s cold; HPC points are squares. **c**, Peak resident memory of the server process against the uncompressed size of the matrix being read (dotted line, equality). On the laptop the server’s peak stays at 82–102 MiB while the layer read grows a hundredfold, from 0.2 to 20 GB. A 160 GB pairwise matrix peaks at 89 MiB on HPC; the idle laptop server uses 79 MiB. **d**, Chunk shape trades cell-row against gene-column reads (one point per shape, cold). Whole-gene chunks make one cell row decompress the whole layer (2.9 s on the laptop at 1 million cells; 7.8 s on HPC); anndata’s default chunks, like the aspect-ratio rule of Procedure Step 10, keep each read to a small part of the layer. Laptop series: AnnZarro 0.4.1 (the v0.4.1 tag, 78c9da6), vectors sent as binary float32. HPC series: the same tag on HPC nodes, cold meaning a fresh server with the node’s page cache evicted.

**Large datasets.** Datasets of tens of millions of cells stay interactive through the subset (**Fig. 7**). We used all 95,624,334 cells of Tahoe-100M ^7^ on a laptop (Apple M3 Max, 128 GiB RAM, headless Chromium 153), stored with raw counts of a 200-gene panel in X and a UMAP ^28^ fitted on 999,073 reference cells sampled evenly across plates and cell lines. Every other cell was placed at the distance-weighted mean of its 15 nearest reference cells in principal-component space, found with faiss ^29^ (recall 0.997 against exact search). On the default 100,000-cell part of all 95.6 million cells, the first plot appeared 4.0 s after the page was opened (3.8 s colored by cell line). Recoloring by another gene then took 0.46 s, stepping to the next of 957 parts 2.8 s, and switching to a subset balanced across the 50 cell lines 3.3 s, including the dialog (**Fig. 7e**; medians over 3 cold runs). These times grow with the number of cells in the dataset, not with the number shown: the first plot took 0.9 s at one million cells. The tab’s renderer process used 0.5 GiB at every dataset size and the server process at most 1.7 GiB.

Every cell can also be drawn at once (**Fig. 7a–c**). In large-plot mode the 95.6 million cells were drawn in 5–6 s at point size 1 and opacity 0.2, colored by cell line or by a gene (3 cold runs each), and zooming in resolves single cells. At the automatic size and opacity the first plot took 6.1–7.1 s colored by a gene and 5.2–5.3 s by cell line (ranges of 3 cold runs); the tab’s renderer process used up to 8.7 GiB and the server, recoloring included, 7.0 GiB (resident memory). On one machine, as in the desktop app, client and server add up: in the tool comparison, which reports physical footprint, drawing every cell with the server on the same laptop took 11.3 GiB of client and 4.6 GiB of server memory, 15.9 GiB together, close to what a 16 GiB laptop offers (**Supplementary Note 1**). Each cell line forms its own island. Colored by *MALAT1*, each island looks uniformly high while the whole dataset is in view, because where cells overlap, large-plot mode draws the highest values on top; zoomed in, each cell shows its own count (median 83 in the region of **Fig. 7c**).

Benchmarks of the server on HPC nodes, the cost of the regular plot, the memory guard’s limits and the largest stores tested are in **Supplementary Note 2**.

## Data availability

The murine bone marrow aging CITE-seq data are available on Zenodo (doi:10.5281/zenodo.15587768; Fred Hutchinson Cancer Center, published 3 June 2025; CC BY 4.0). Tahoe-100M is available from Hugging Face (tahoebio/Tahoe-100M, CC0 1.0) ^7^. The *C. elegans* store of **Fig. 4d** is not deposited; the script docs/_tools/datasets/celegans_connectome.py in the AnnZarro repository (v0.4.1) builds it from public downloads, checked by sha256 (https://annzarro.readthedocs.io/en/latest/paper/cell-by-cell.html#cell-by-cell-connectome): the wiring diagram of Varshney et al. ^14^ (WormAtlas NeuronConnect.xls; article CC BY), CeNGEN neuron-type expression ^15^ (cengenDataSC, GPL-3.0) and neurotransmitter identities from Wang et al. ^30^ (CC BY 4.0). The processed AnnZarro-ready store, bm_aging_annzarro.zarr (derived from doi:10.5281/zenodo.15587768, CC BY 4.0), and the Tahoe-100M UMAP of every cell with its list of reference cells are deposited on Zenodo (doi:10.5281/zenodo.23277103) ^31^. The Visium store of **Fig. 2d** is built from the 10x Genomics dataset by docs/_tools/make_spatial_demo.py in the AnnZarro repository (v0.4.1). The measurements behind **Figs. 1c, 7 and 9**, the benchmark code and an evidence site that links every number in this paper to its source are in the repository https://github.com/settylab/annzarro-reproducibility, which will be made public with this preprint.

## Code availability

AnnZarro is released under the MIT license at https://github.com/settylab/annzarro, on PyPI as annzarro, and as standalone desktop applications on GitHub releases; this protocol describes version 0.4.1 (tag v0.4.1, commit 78c9da6; Zenodo doi:10.5281/zenodo.23222217). The script for data preparation (Steps 3–10, docs/_tools/datasets/bm_aging.py) and the panel sets for the worked examples (docs/_static/panelsets/protocol) are in the AnnZarro repository. The benchmark code and the scripts behind the figures are in https://github.com/settylab/annzarro-reproducibility (see Data availability). Documentation is at https://annzarro.readthedocs.io. Every figure of this paper can be reproduced from it: a page per figure (https://annzarro.readthedocs.io/en/latest/paper/) links to the share links and panel sets that reopen each view, and its tutorials rebuild **Figs. 2 and 4–6** step by step in the app.

## Acknowledgments

We thank members of the Setty laboratory for discussions and for testing AnnZarro. This work was supported by the National Institute of General Medical Sciences of the National Institutes of Health under award numbers R35GM147125 (M.S.) and K99GM159044 (D.J.O.). This work used Fred Hutch Scientific Computing resources supported by NIH grants S10OD038189 and S10OD028685.

## Use of AI assistance

The authors used AI coding agents (Claude, Anthropic, through Claude Code) to write and test software and analysis code, run and analyze the benchmarks, audit the compared tools, and draft and edit the manuscript. D.J.O. and S.B. tested and edited the code by hand, and D.J.O. checked the full text and every figure against the running software and the underlying data; no figure contains AI-generated imagery. The source of each number and comparison mark in the paper was recorded and checked against the text. The authors take full responsibility for the content.

## Author contributions

D.J.O. conceived and developed AnnZarro and wrote the manuscript. S.B. implemented h5ad support and contributed to the user interface. M.S. supervised the work and edited the manuscript.

## References

[1] Palla, G. et al. Squidpy: a scalable framework for spatial omics analysis. Nat. Methods, 19, 171–178, 2022. doi: 10.1038/s41592-021-01358-2.

[2] Cang, Z. et al. Screening cell–cell communication in spatial transcriptomics via collective optimal transport. Nat. Methods, 20, 218–228, 2023. doi: 10.1038/s41592-022-01728-4.

[3] The MICrONS Consortium,. Functional connectomics spanning multiple areas of mouse visual cortex. Nature, 640, 435–447, 2025. doi: 10.1038/s41586-025-08790-w.

[4] Gamlin, C. R. et al. Connectomics of predicted Sst transcriptomic types in mouse visual cortex. Nature, 640, 497–505, 2025. doi: 10.1038/s41586-025-08805-6.

[5] Virshup, I., Rybakov, S., Theis, F. J., Angerer, P. & Wolf, F. A. anndata: Access and store annotated data matrices. J. Open Source Softw., 9, 4371, 2024. doi: 10.21105/joss.04371.

[6] Otto, D. J., et al. Comparing phenotypic manifolds with Kompot: cluster-free differential expression at single-cell resolution. bioRxiv, 2025. doi: 10.1101/2025.06.03.657769.

[7] Zhang, J. et al. Tahoe-100M: mapping drug-induced molecular phenotypes at single-cell resolution. Cell, 189, 5945–5961.e8, 2026. doi: 10.1016/j.cell.2026.08.035.

[8] Setty, M. et al. Characterization of cell fate probabilities in single-cell data with Palantir. Nat. Biotechnol., 37, 451–460, 2019. doi: 10.1038/s41587-019-0068-4.

[9] Otto, D. J., Jordan, C., Dury, B., Dien, C. & Setty, M. Quantifying cell-state densities in single-cell phenotypic landscapes using Mellon. Nat. Methods, 21, 1185–1195, 2024. doi: 10.1038/s41592-024-02302-w.

[10] Rue-Albrecht, K., Marini, F., Soneson, C. & Lun, A. T. L. iSEE: Interactive SummarizedExperiment explorer. F1000Res., 7, 741, 2018. doi: 10.12688/f1000research.14966.1.

[11] Keller, M. S. et al. Vitessce: integrative visualization of multimodal and spatially resolved single-cell data. Nat. Methods, 22, 63–67, 2025. doi: 10.1038/s41592-024-02436-x.

[12] 10x Genomics, Mouse brain serial section 1 (sagittal-anterior), spatial gene expression dataset analyzed using Space Ranger 1.1.0. https://www.10xgenomics.com/datasets/mouse-brain-serial-section-1-sagittal-anterior-1-standard-1-1-0, 2020. License CC BY 4.0. Files: https://cf.10xgenomics.com/samples/spatial-exp/1.1.0/V1_Mouse_Brain_Sagittal_Anterior/. Accessed 2026-10-02.

[13] Persad, S. et al. SEACells infers transcriptional and epigenomic cellular states from single-cell genomics data. Nat. Biotechnol., 41, 1746–1757, 2023. doi: 10.1038/s41587-023-01716-9.

[14] Varshney, L. R., Chen, B. L., Paniagua, E., Hall, D. H. & Chklovskii, D. B. Structural properties of the *Caenorhabditis elegans* neuronal network. PLoS Comput. Biol., 7, e1001066, 2011. doi: 10.1371/journal.pcbi.1001066.

[15] Taylor, S. R. et al. Molecular topography of an entire nervous system. Cell, 184, 4329–4347.e23, 2021. doi: 10.1016/j.cell.2021.06.023.

[16] Megill, C. et al. cellxgene: a performant, scalable exploration platform for high dimensional sparse matrices. bioRxiv, 2021. doi: 10.1101/2021.04.05.438318.

[17] CZI Cell Science Program, et al. CZ CELLxGENE Discover: a single-cell data platform for scalable exploration, analysis and modeling of aggregated data. Nucleic Acids Res., 53, D886–D900, 2025. doi: 10.1093/nar/gkae1142.

[18] Speir, M. L. et al. UCSC Cell Browser: visualize your single-cell data. Bioinformatics, 37, 4578–4580, 2021. doi: 10.1093/bioinformatics/btab503.

[19] Tarhan, L., et al. Single cell portal: an interactive home for single-cell genomics data. bioRxiv, 2023. doi: 10.1101/2023.07.13.548886.

[20] Li, B. et al. Cumulus provides cloud-based data analysis for large-scale single-cell and single-nucleus RNA-seq. Nat. Methods, 17, 793–798, 2020. doi: 10.1038/s41592-020-0905-x.

[21] Davie, K. et al. A single-cell transcriptome atlas of the aging *Drosophila* brain. Cell, 174, 982–998.e20, 2018. doi: 10.1016/j.cell.2018.05.057.

[22] Ouyang, J. F. et al. ShinyCell: simple and sharable visualization of single-cell gene expression data. Bioinformatics, 37, 3374–3376, 2021. doi: 10.1093/bioinformatics/btab209.

[23] Chen, B. J., et al. ShinyCell2: an extended library for simple and sharable visualisation of spatial, peak-based and multi-omic single-cell data. bioRxiv, 2025. doi: 10.1101/2025.04.22.650045.

[24] Lun, A. T. L. & Kancherla, J. Powering single-cell analyses in the browser with WebAssembly. J. Open Source Softw., 8, 5603, 2023. doi: 10.21105/joss.05603.

[25] Vieth, A. et al. ManiVault: a flexible and extensible visual analytics framework for high-dimensional data. IEEE Trans. Vis. Comput. Graph., 2023. doi: 10.1109/TVCG.2023.3326582.

[26] Chari, T. & Pachter, L. The specious art of single-cell genomics. PLoS Comput. Biol., 19, e1011288, 2023. doi: 10.1371/journal.pcbi.1011288.

[27] Fred Hutch Cancer Center,. Single-cell CITE-seq of murine bone marrow across aging (Young, Mid, Old). Zenodo, 2025.

[28] McInnes, L., Healy, J., Saul, N. & Großberger, L. UMAP: Uniform manifold approximation and projection. J. Open Source Softw., 3, 861, 2018. doi: 10.21105/joss.00861.

[29] Johnson, J., Douze, M. & Jégou, H. Billion-scale similarity search with GPUs. IEEE Trans. Big Data, 7, 535–547, 2021. doi: 10.1109/TBDATA.2019.2921572.

[30] Wang, C. et al. A neurotransmitter atlas of *C. elegans* males and hermaphrodites. eLife, 13, RP95402, 2024. doi: 10.7554/eLife.95402.

[31] Otto, D. J., Baasri, S. & Setty, M. AnnZarro paper data: an AnnZarro-ready store of murine bone marrow across aging, and a UMAP of every cell of Tahoe-100M. Zenodo, 2026. Version 1.0.

