## Supplementary Notes for "AnnZarro: scalable, interactive exploration of cell-by-cell and gene-by-gene relationships in single-cell data"

### Supplementary Note 1: Comparison of interactive viewers

The dataset sizes (open diamonds) are the largest single datasets each project documents, opened in full or on a subset, or, for CELLxGENE and UCSC, the largest observed on their public portals and drawn in full in a browser. Where a source gives only a lower bound ("more than"), the diamond marks that bound. They demonstrate a size and do not mark a limit. Each project's documentation and source code were checked between 3 and 6 October 2026. A tool that downloads a whole file before loading it does not read a remote store. Each tool was also checked in its running application where it could be run. "Cell × cell matrix as color or axis" is full only when the user can tell which cell the shown row belongs to (for example by focusing that cell); a row picked by index scores half. The focus column is strict: a focused cell or gene must re-drive every linked panel through its stored pairwise row; a gene color that reaches several views does not count. "Lazy chunked reads" and "every cell drawn" score what the GUI can do, also after a user choice or a preparation step; where it takes writing code, the mark is half. "User picks a subset of cells" is full only when the tool then loads and draws only those cells; a subset that hides or grays the other cells scores half. "Automatic subset for large data" is full when a dataset too large to draw whole opens by default on a subset of its cells, the rest still reachable. Only the core tool is scored, not its extension packages.

**Measured glyphs (cells drawn within a 16 GiB client memory budget).** Each tool opened datasets built from Tahoe-100M, with a 200-gene panel in X (about 110 stored values per cell against about 1,470 in the full transcriptome): its first *N* cells (from the store used for **Fig. 7**) and, beyond its full size, stores that repeat its cells (below). Each drew a UMAP colored by cell line and by *MALAT1* (metastasis associated lung adenocarcinoma transcript 1), while the physical footprint of its processes (macOS phys_footprint, what Activity Monitor calls Memory) was sampled on a laptop (Apple M3 Max, 128 GiB RAM, macOS). The value is the largest *N* whose client peak stayed within 16 GiB: all browser processes for a web tool, whose server is treated as remote, and the whole R or Python process for Scanpy and Seurat; the server peak was measured in the same run. iSEE and ShinyCell2 render on the server, so for them the server is reported. Versions: AnnZarro 0.4.1, CELLxGENE Annotate 1.3.0, Cirrocumulus 1.1.61, UCSC Cell Browser 1.2.20, iSEE 2.18.0, ShinyCell 2.1.0, Scanpy 1.12.4, Seurat 5.5.1, Vitessce as hosted at vitessce.io, Kana 3.2.0 as served at kanaverse.org, ManiVault Studio 1.5.0-dev, Loupe Browser 9.1.0 (Electron 28.3.2, Chromium 120), and headless Chromium 153 (driven by Playwright 1.63) for the web tools. Tahoe-100M has 95.6 million cells; stores above that are synthetic, built for these benchmarks. For this column, cell *i* of a larger store copies Tahoe-100M cell *i* modulo its number of cells, with its obs, its expression values and its embedding coordinates plus a small random jitter, so stored values per cell and gene distributions are Tahoe's, while every demonstration plot and screenshot uses real data. The figure draws one glyph, the largest size a tool drew in full; what stopped it at the next size of the ladder is stated here. For Vitessce, Scanpy and Seurat the client's memory passed the budget. Kana drew 1 million cells; at 2 million its WebAssembly module aborted in the browser. SCope drew 1 million cells; at 2 million its browser renderer crashed in both attempts, at 4.7 GiB of client memory, far below the budget. Each tool's input was prepared in the format it needs (a zarr store, an h5ad file, a built site, an app, a SingleCellExperiment); preparing it, by conversion or build, is not measured for any tool, AnnZarro included. Servers have a budget too, a flat 32 GiB of physical footprint (the concurrent peak of the process tree), applied to every tool, AnnZarro included; a tool whose server passed it stopped at that size. CELLxGENE Annotate drew 14.1 million cells within it and its server reached 35.9 GiB at 16.8 million (34.4 GiB in `--backed` mode, which also stopped at 14.1 million); Cirrocumulus drew 10 million, its server reaching 43.6 GiB at 14.1 million. AnnZarro's server stayed within it at every size; its concurrent peak, 8.5 GiB, came with every cell of the 200-million-cell store. Other tools failed at the next size within the budget: they crashed, as Kana's WebAssembly module and the renderers of SCope and Loupe Browser did, or could not read their input. UCSC drew every cell of Tahoe-100M, 95.6 million, the largest real dataset, where its ladder stops. ShinyCell2 drew 10 million cells, its server at 7.2 GiB (the HPC's process-tree resident memory); at 20 million its app builder, makeShinyApp, refuses the HDF5-backed input, which holds more than $2^{31}-1$ values, an input path limit. Cirrocumulus's prepared path drew 10 million cells; at 20 million its server passed the budget (33.9 GiB, the HPC's process-tree resident memory, the only measure there). iSEE, measured on its HDF5-backed SingleCellExperiment, drew 10 million cells with its server at 24.6 GiB; at 20 million its server passed the budget (49.5 GiB). Seurat's default in-memory matrix (dgCMatrix) holds at most $2^{31}-1$ values, about 19.2 million cells on this panel, but reaches the 16 GiB budget at 2 million cells, long before that, so the limit sets no glyph; the BPCells variant keeps the matrix on disk and avoids it. An arrow marks a lower bound, the largest size tested. A run counts as hung when client and server together use less than 2 CPU-seconds over 300 s; a size with at least 3 hung attempts is a client stop. AnnZarro drew 175 million cells within the budget and every cell up to 200 million, where its client used 17.4 GiB; at 250 million cells its memory guard declined the first plot ("250,000,000 points is above 200,000,000, the largest plot tested to draw", the guard's own message). Rings are measured for the tools whose subset reduces what is loaded ("user picks a subset" full, or "automatic subset" full or half): the largest store each opened and drew with its subset within the same client budget. With a subset, AnnZarro opened and drew every store up to the largest tested, 1.5 billion cells, so 1.5 billion is a lower bound, not a limit. This held with its default subset (100,000 cells shown above 200,000) and with a 1,000,000-cell subset, and the client used at most 1.5 GiB and 2.4 GiB, respectively, at any size. Server memory at 1 billion cells is given two ways. Summed over the server's processes, each at its own maximum during the first plot, it was 4.0 GiB; at its highest moment, all processes together used 3.9 GiB, the value the 32 GiB budget limits. With the server on the same machine, as in the desktop app, a user needs the second, and the first bounds it from above. The first plot of a 1-billion-cell store took 39 s with the default subset and 41 s with the 1,000,000-cell subset. Most of that time (27 s) went to listing the subset's cell names; each UMAP column took 3 s to its first byte. Single Cell Portal is hosted only and shows its documented size. The macOS budget was measured, not enforced; under a hard 16 GiB limit in Docker (Linux), Scanpy and Seurat agreed with the measured crossings, except that Scanpy also fitted 8.4 million cells, so its 7.1 million is conservative by one step. AnnZarro was measured with its server on the same laptop, as in the desktop app, and its value counts the client; drawing all 95.6 million cells, recoloring included, took 11.3 GiB of client and 4.6 GiB of server memory. Three tools were not measured, for these reasons. CELLxGENE Explorer and Single Cell Portal are hosted services that cannot load our data. VizIt draws a sample of 50,000 points fixed when the data are prepared, so it never draws every cell. ManiVault Studio and Loupe Browser, desktop applications with no scripted route into them, were measured by hand in an interactive desktop session on the same inputs.

- **AnnZarro.** 1.5 billion cells opened and drawn on a subset, the largest store tested and so a lower bound (synthetic stores, this paper's measurement above); 95,624,334 cells of Tahoe-100M, every cell drawn in one plot on request (**Fig. 7**). Data are read in chunks on demand on the server; above 200,000 cells the default view is a reproducible 100,000-cell subset, whose parts reach every cell, and one setting draws every cell in one plot.
- **CELLxGENE.** Two programs, scored as two rows: Explorer, the hosted viewer of CELLxGENE Discover, and Annotate, the self-hosted package, whose README claims "at least 1 million cells". Annotate accepts an S3, GCS or HTTP URL but downloads the whole file before loading it, which does not count as reading a remote store, and its documentation cautions about multi-user hosting without documenting a multi-user deployment, so "hosted multi-user" scores half. "Lazy chunked reads" scores half: with `--backed`, Annotate reads genes lazily only when X is stored column-major (CSC), and differential expression only when it is row-major (CSR), never both; on an 8,090-cell matrix, each gene request took 0.53 s and added 203 MiB of server memory under the row-major layout scanpy writes by default, against 0.02 s and 1 MiB column-major. Explorer: 11,441,407 cells, the mouse embryo timelapse of Qiu et al. 2024 on CELLxGENE Discover (`https://cellxgene.cziscience.com/e/dcfd4feb-18a3-4b30-81d7-1b0c544a8ab3.cxg/`); in three headless-Chromium runs on 4 October 2026 the hosted Explorer drew every cell in one draw call, the first full render taking 10.8–17.1 s and the page reporting "11441407 of 11441407 cells". Discover's documentation states a lower figure, 4.3 million cells, as a submission and hosting policy; the dataset above shows that the Explorer draws more: "CELLxGENE Discover sets the maximum per dataset file size for submissions to 50 GB. Additionally, datasets with more than 4.3 million cells can be submitted but will not visualized [sic] in CELLxGENE Explorer" (`https://github.com/chanzuckerberg/single-cell-data-portal/blob/main/frontend/doc-site/032__Contribute%20and%20Publish%20Data.mdx`, line 31, section Scale Constraints; also at commit 9659d41). The Discover paper gives the same figure as a performance statement: Explorer explores "individual datasets for up to 4.3 million cells in <1 min" (Nucleic Acids Res. 53, D886, 2025). Every cell is drawn (WebGL draw count = number of cells); the server reads TileDB slices on demand; "Subset to currently selected cells".
- **Vitessce.** 1,283,972 cells (Salcher et al. 2022 lung cancer atlas), a public demo of the project. Every cell drawn (one deck.gl scatterplot layer over the full embedding); genes read by HTTP range from chunked zarr in the browser; cell sets highlight but do not subset.
- **Cirrocumulus.** 1.3 million cells (documentation, "Primary embedding"). Every cell drawn; prepared data are read partially over the network, while `cirro launch` on an h5ad holds it in server memory; selections dim or shrink the other cells, and setting the opacity of filtered-out cells to 0 shows only the selection, a view-only subset made with a styling slider rather than a labeled command (half under the subset rule above).
- **iSEE.** More than 170,000 cells, a mass cytometry study of PBMCs in the iSEE paper (Rue-Albrecht et al. 2018, "Mass cytometry" use case); the largest single-cell RNA-seq set in its documentation is 68,579 cells (10x PBMC 68k, bigdata vignette). Every cell drawn by default, optional grid downsampling per panel; HDF5-backed assays read on demand. The extension package iSEEu adds panels such as on-the-fly marker tables; extensions are not scored. Plots are drawn on the server (server-rendered in **Fig. 1c**).
- **UCSC Cell Browser.** 21,786,931 cells, PanSci Organismal Aging "All" (`https://cells.ucsc.edu/?ds=mouse-pansci+all`; sampleCount in the portal's dataset.json); in three headless-Chromium runs on 4 October 2026 all 21,786,931 points were drawn in one draw call, the first render taking 7.3–7.4 s. The documentation states only a portal total ("over 160 million cells" across 1,600 datasets). Every cell drawn; genes fetched by HTTP byte range from static files; "Only show selected".
- **ShinyCell2.** 161,764 cells in the ShinyCell2 CITE-seq tutorial. The row scores ShinyCell2, which adds

spatial coordinates and image underlays to ShinyCell; ShinyCell advises about 30,000 cells per 8 GiB of RAM when building an app. Every cell drawn; one HDF5 chunk per gene read on demand; per-plot "Toggle to subset cells". Plots are drawn on the server (server-rendered).

- **Single Cell Portal.** 3,177,310 cells, study SCP2805 (human retina, snRNA-seq), the largest study with an embedding on the portal, read from the portal's public API. Above 100,000 cells the default view is a stratified 100,000-cell sample, and "All Cells" in the menu draws every cell; expression is read per gene on the server.
- **Kana.** 173,954 cells (Zilionis et al. 2019 lung data, Kana's paper). Every cell is drawn, and the whole matrix is held in the browser's WebAssembly memory; a user can analyze a subset chosen at load time.
- **SCope** 1.8.2, as served at scope.aertslab.org. The measured copy was built locally from 1.8.2 (9ae5207) with nine build-environment changes and run with four run-time settings, among them its frontend served by a static file server and its database migration step skipped; SCope's code is unchanged. 567,950 cells drawn in the Fly Cell Atlas session (whole-atlas 10x "Relaxed" loom), counted from the viewer's state. Every cell is drawn, and the server reads one gene row from the loom file per request. Its last desktop build is a 2019 prerelease that its maintainers describe as likely not to work, so the desktop column scores no.
- **Loupe Browser** 9.1.0, checked in the running app on the demo data; spatial and image underlays from its documentation, lazy reads measured in the app (below). Release notes mention datasets of more than 1 million cells; how points are drawn and loaded is not documented. Measured in an interactive desktop session on the same inputs (converted to .cloupe with LoupeR 1.1.5), it drew every cell up to 5 million within the client budget, at 5.7 GiB; at 10 million it drew the cell lines, but the window went blank when coloring by a gene, because the renderer process exited at 4.0 GiB, well under the budget; a per-process memory limit of its embedded Chromium would explain this, but was not established. Lazy reads were measured in the app on two .cloupe files with the same 1,000,000 cells and 200 or 2,000 genes (0.37 and 2.20 GiB), each opened three times in a freshly started app. The memory after opening did not depend on the file, a median of 1.27 GiB for both, with a largest spread of 0.185 GiB between repetitions of one file, so Loupe does not load the expression matrix when it opens a file. Coloring by *MALAT1* raised the footprint by 0.1 to 0.3 GiB, and the peak while loading was 2.0 to 2.2 GiB, for both files. This shows that the matrix is not loaded whole at 1 million cells; how Loupe reads the file is not documented and was not established. Loupe reported 999,966 barcodes, not 1,000,000, and 34 cells have no *MALAT1* count in the inputs, so its legend appears to count the cells with a nonzero value of the colored gene (tested for this one gene).
- **ManiVault Studio** 1.5.0-dev (desktop; the official 1.5.0 macOS installer reports itself as 1.5.0-dev because it was built from a pre-release commit, and no release-tagged macOS build exists); Cytosplore Viewer 5.0.0, a separate build on the same framework, was not examined. A lasso selection becomes a named cluster that is exported with its cells (annotation editing). It loads obsp by default and colors an embedding by a stored obsp row, picked by its row number rather than by a focused cell, so the user cannot tell which cell the row belongs to (half under the rule above); varp is not read. The documented largest dataset is 15 million cells, a mass cytometry (CyTOF) dataset in the HSNE paper[1]. Measured in an interactive desktop session on the same inputs, written with anndata 0.11.4 (ManiVault 1.5.0 does not open h5ad files written by anndata 0.13 with pandas 3, which store the index as a group), it drew every cell up to 1 million within the client budget, at 12.7 GiB, and needed 24.4 GiB at 2 million.
- **VizIt** 1.0.0, as deployed on pd5d.yale.edu. 309,960 cells observed on its hosted PD5D atlas, of which the UMAP draws 50,000: a random sample fixed in code at preparation, so "every cell drawn" is not found and "automatic subset" scores half; expression is fetched one gene per request; choosing samples limits what is drawn, not what is loaded.
- **Scanpy and Seurat**, the baseline of what code can do. 1.3 million cells (Scanpy paper; Seurat v5

sketch vignette); Scanpy’s experimental dask tutorial plots about 1.4 million cells. Every cell drawn by default (Seurat rasterizes above 100,000 cells, without dropping any); the whole matrix is in memory unless backed or lazy modes are chosen in code; subsets by code.

### Supplementary Note 2: performance benchmarks

Moved from Anticipated results, which now leads with what a user sees. The laptop series and the chunk shapes are in **Fig. 9**; the Tahoe-100M series are in **Fig. 7**.

Memory stays small on the server as well as in the browser: on the laptop, the server's peak resident memory while serving a gene column or a cell row stayed at 82–102 MiB for layers of 0.2 to 20 GB uncompressed, against 79 MiB idle (**Fig. 9c**).

The Tahoe-100M ladder of **Fig. 7** was repeated on an HPC node (36 CPUs, 755 GiB RAM; stores on /hpc/temp, an NFSv4.1 scratch filer; a fresh server and the node's page cache evicted before each cold request): with AnnZarro v0.4.1 the server returned the gene column of X for every cell in 0.16 s cold at 1 million cells and 2.8 s at 95.6 million, and in 2.2 s at 95.6 million for a gene detected in fewer cells (medians of 3 cold requests).

Controlled benchmarks on HPC nodes (Xeon Gold 6254, a whole 36-CPU node, one server process; data on /hpc/temp, an NFSv4.1 scratch filer; a fresh server and the node's page cache evicted before each cold request) repeat these measurements on larger matrices and larger stores, with AnnZarro v0.4.1. On one node, one row of a dense cells × cells Gaussian kernel took 0.29, 0.31 and 0.30 s cold for 50,000, 100,000 and 200,000 cells, and a row of a dense diffusion-distance matrix 0.31, 0.29 and 0.27 s (**Fig. 9b**); the largest matrix is 160 GB uncompressed. Peak server memory while serving such a row stayed at 85–89 MiB across distance and correlation matrices of 1.6–160 GB, against 78 MiB idle. A gene-by-gene correlation row for 30,000 genes took 0.24 s. On a 1.17-million-cell prefix of Tahoe-100M, a gene column of X took 0.20 s cold and 0.01 s warm and a kNN row 0.21 s cold. Chunk shape decides, by construction, what a cell-row read costs: with whole-gene chunks every chunk holds part of each cell's row, so one row decompresses the whole layer, whereas in the Step 10 shape, 8192 × 64, it decompresses only the chunks of its own block of cells. One cell row of a 1-million-cell, 5,000-gene layer of Tahoe-100M with whole-gene chunks took 7.8 s cold, and 0.38 s after rechunking.

The regular plot keeps hover and focus, and it is the costly one. At 1 million cells, the most it draws, its first plot colored by a gene took 2.7 s and filled up to 0.9 GiB of the tab's 4.1 GiB JavaScript heap; in large-plot mode, 2 million cells took 0.87 s and 5 million 1.1 s (**Fig. 7f,g**; 3 cold runs each). Large-plot mode therefore starts above 1 million points, so that several regular plots fit side by side.

With the memory guard at its default settings, every cell was drawn for stores up to 200 million cells (**Supplementary Note 1**); at 250 million cells the guard declined the first plot before anything was drawn, with the message "250,000,000 points is above 200,000,000, the largest plot tested to draw". Large-plot mode keeps the points' positions off the JavaScript heap (0.02 GiB of heap at 95.6 million cells), and Chromium caps the tab's heap at 4.1 GiB whatever the browser flags (**Fig. 7g**); the positions are held in the tab's renderer process instead, whose memory grows with the points drawn. With a subset of 100,000 cells (the default) or 1,000,000, every store tested opened and drew, up to the largest at 1.5 billion cells, with at most 2.4 GiB of client memory (physical footprint). At 1 billion cells the server's processes, each counted at its own peak, used 4.0 GiB together, and the first plot took 39–41 s, most of it spent listing the subset's cell names (**Supplementary Note 1**). Without the guard, at 250 million cells the tab could not allocate the buffer for the positions, and the panel reported the error instead of drawing.

### References


[1] van Unen, V. et al. Visual analysis of mass cytometry data by hierarchical stochastic neighbour embedding reveals rare cell types. *Nat. Commun.*, **8**, 1740, 2017. doi: 10.1038/s41467-017-01689-9.
